# Systemically targeting monocytic MDSCs using dendrimers and their cell-level biodistribution kinetics

**DOI:** 10.1101/2024.02.14.580395

**Authors:** Chad A. Littrell, Gregory P. Takacs, Chenikkayala Siva Sankara, Alexandra Sherman, Kai A. Rubach, Julia S. Garcia, Coral Bell, Jeffrey K. Harrison, Fan Zhang

## Abstract

The focus of nanoparticles *in vivo* trafficking has been mostly on their tissue-level biodistribution and clearance. Recent progress in the nanomedicine field suggests that the targeting of nanoparticles to immune cells can be used to modulate the immune response and enhance therapeutic delivery to the diseased tissue. In the presence of tumor lesions, monocytic-myeloid-derived suppressor cells (M-MDSCs) expand significantly in the bone marrow, egress into peripheral blood, and traffic to the solid tumor, where they help maintain an immuno-suppressive tumor microenvironment. In this study, we investigated the interaction between PAMAM dendrimers and M-MDSCs in two murine models of glioblastoma, by examining the cell-level biodistribution kinetics of the systemically injected dendrimers. We found that M-MDSCs in the tumor and lymphoid organs can efficiently endocytose hydroxyl dendrimers. Interestingly, the trafficking of M-MDSCs from the bone marrow to the tumor contributed to the deposition of hydroxyl dendrimers in the tumor. M-MDSCs showed different capacities of endocytosing dendrimers of different functionalities *in vivo*. This differential uptake was mediated by the unique serum proteins associated with each dendrimer surface functionality. The results of this study set up the framework for developing dendrimer-based immunotherapy to target M-MDSCs for cancer treatment.

## Introduction

Nanoparticles have been extensively used to deliver therapeutic payloads for disease treatment. While most *in vivo* biodistribution studies of nanoparticles have been focused on their tissue-level accumulation and clearance^1, 2^, recent progress in the nanomedicine field suggested that targeting nanoparticles to immune cells can be used to modulate the immune response and to enhance therapeutic delivery to the disease region^3–5^.

For example, monocytic-myeloid-derived suppressor cells (M-MDSCs) are important cellular targets in cancer^6^. M-MDSCs are pathologically activated immature monocytes with potent immunosuppressive activities. Clinically, a high burden of M-MDSCs is associated with poor prognosis of many solid tumors^7^. In cancer, M-MDSCs help create and maintain an immunosuppressive tumor microenvironment (TME)^8, 9^. These cells can suppress anti-tumor T cells and promote regulatory T cells and anti-inflammatory myeloid cells^8, 9^. As such, there is an urgent need to develop delivery strategies to target M-MDSCs for cancer treatment.

In addition to their immunosuppressive features, M-MDSCs, as phagocytes^10, 11^, could also significantly affect the *in vivo* fate of nanoparticles. Historic studies have established the critical roles of myeloid cells in clearing nanoparticles^12, 13^, while emerging evidence has shown that myeloid cells in circulation can take up nanoparticles and actively transport them to the inflamed tissue^4, 14, 15^. M-MDSCs are significantly elevated in the peripheral blood of high-grade glioblastoma patients, accounting for as much as 10% of total cells in the peripheral blood and 30% of total peripheral blood mononuclear cells^16, 17^. In the presence of tumor lesions, the bone marrow accelerates monopoiesis and enhances the egress of M-MDSCs to the systemic circulation, leading to significant expansion of their population in the peripheral blood and in the spleen^18^. Tumors constantly recruit M-MDSCs in large amounts through CCR2-mediated chemotaxis to replenish the tumor-associated macrophages (TAMs)^19, 20^. The abundance of M-MDSCs and their constant infiltration to the tumor sites highlights the potential of M-MDSCs in mediating nanoparticle deposition at the tumors.

Given the important roles of M-MDSCs in establishing the TME and mediating nanoparticle tumor-targeting, many efforts have tried to establish the correlation between nanoparticle physiochemical properties, such as size, surface charge, and surface ligands to their targeting of M-MDSCs^21–24^. However, a critical challenge in studying the cell targeting behaviors of nanoparticles is that when nanoparticles are injected into the blood, multiple serum proteins such as immunoglobulins, fibrinogen, complement proteins, and apolipoproteins readily adsorb to the nanoparticle surface, forming a ‘protein corona’^25, 26^. The protein corona masks the native nanoparticle interactions with the cell surface and alters the nanoparticle’s cellular tropism^25, 26^. It is now recognized that nanoparticle-associated proteins dictate nanoparticle interactions with cells and, more broadly, their *in vivo* targeting behaviors. The physical properties of nanoparticles only play a secondary role in this process.

Dendrimers represent a class of ultra-small nanoparticles with sub-10nm size that carry drug payload on their surface. About 26 dendrimer-based therapeutics with various types of payloads are currently under Phase I-III clinical trials^27^. Previous studies showed that systemically administrated hydroxyl-terminated PAMAM dendrimer (OH dendrimer) can selectively target TAMs in murine glioblastoma (glioma) models^28, 29^. However, given the heterogeneous nature of TAMs, it is unclear what subset(s) are being targeted and what mechanism mediates the selective cell-targeting. In this study, we quantitatively examined the cell-level biodistribution kinetics after systemically administrating OH dendrimers in two murine models of glioma. M-MDSCs can efficiently endocytose OH dendrimers in all reservoir tissues (bone marrow, spleen, peripheral blood, and tumor). In the tumor, M-MDSCs and microglia showed a high capacity of taking up OH dendrimers and these two cellular compartments accounted for more than half the amount of the OH dendrimer deposition in the tumor. The trafficking of M-MDSCs from bone marrow to the tumor region contributed to the tumor deposition of OH dendrimers. We also found the surface functionality of dendrimers significantly affected their ability to ‘target’ M-MDSCs. Finally, the unique serum proteins associated with each dendrimer surface functionality affected the dendrimer’s differential target to M-MDSCs.

## Materials and Methods

### Generation of Cy5-labeled dendrimers

The G6 PAMAM hydroxyl (OH), amine (NH_2_), and succinamic acid (SA) surface dendrimers were purchased from (Dendritech, Inc). The NH_2_ surface dendrimer was used for reaction after the evaporation of methanol from the stock solution. The OH and SA surface dendrimers were further functionalized with amine terminals to conjugate Cy5 mono NHS ester (Cytiva). For the Cy5-labeling of OH dendrimers: Step 1: Fmoc-GABA-OH (Sigma-Aldrich) was coupled with G6-OH using PyBOP (Merck) as a coupling reagent to produce an intermediate with protecting group Fmoc. Step 2: The Fmoc protecting group was removed using piperidine (Sigma-Aldrich) – DMF (Sigma-Aldrich) mixture to produce bi-functional dendrimers. The crude was purified by dialyzing (membrane cutoff = 12–14 kDa) against DMF (Sigma-Aldrich) for 24 h by changing the DMF every 8 h. Step 3: The conjugation of Cy5 mono NHS ester was carried out in the presence of borate buffer (pH 8.5) with pure bifunctional dendrimer to produce G6-OH-Cy5 conjugate. Similarly, amine surface G6 dendrimer was labeled with Cy5 using borate buffer (pH 8.5). For the Cy5-labeling of SA dendrimers, in step 1, G6 succinamic acid surface dendrimer was coupled with N-Fmoc 1,5- diaminobutane hydrobromide (Sigma-Aldrich) using EDC.HCl (Sigma-Aldrich) as a coupling reagent. In step 2, the successful deprotection of Fmoc using piperidine (Sigma-Aldrich) DMF (Sigma-Aldrich) mixture resulted in bi-functional dendrimers. The Cy5-labeling of bi-functional dendrimers under DMSO (Sigma-Aldrich) and DIEA (Sigma-Aldrich) produced G6-succinamic acid-Cy5 conjugate in step 3. The synthesized G6 PAMAM-Cy5 conjugates are in good agreement with the reported literature data^30^.

### Cell lines

The GL261 glioma cells were cultured in in RPMI (Invitrogen) with 1% penicillin-streptomycin (Invitrogen), 10% fetal bovine serum (FBS) (Thermo Scientific), and 4mM L-Glutamine. KR158 glioma cells were maintained in Dulbecco modified Eagle medium (DMEM) supplemented with 10% heat-inactivated fetal bovine serum (FBS) and 1% penicillin–streptomycin. Cell cultured in wells were expanded in T75 flasks (Falcon) in a humidified incubator (Thermo Scientific) at 37°C with 5% CO_2_. All cell lines tested negative for mycoplasma based on DNA-based PCR tests.

### M-MDSC induction and culture

Induction of M-MDSCs from transgenic CCR2^RFP/WT^/CX3CR1^GFP/WT^ bone marrow cells or WT C57BL/6 was adapted from previously published work using wildtype C57BL/6 mice^31^. Bone marrow cells collected from the femur were seeded at a density of 1 x 10^6^ cells/mL in KR158 cell-conditioned culture media (50% v/v KR158 conditioned media + 50% RPMI-1640 (Gibco) + 10% FBS (Corning) + 1% penicillin-streptomycin (Corning), + 1% GlutaMax (Gibco) + 1% Non-Essential Amino Acids (Gibco), 0.22µm sterile bottle-top filtered). Cells were cultured for five days. At the endpoint, suspension cells were collected from the supernatant and adherent cells by scraper (Fisher Scientific) after 15-minute incubation at 37°C, 5% CO_2_ with enzyme-free cell dissociation buffer (Gibco). Flasks were twice-washed using 10-25mL FACS buffer (10% FBS + 1× HBSS) and all cells were collected by centrifugation (500 x g for 5 minutes at 4°C). Cells were collectively resuspended in a 50mL sterile conical (Falcon) in FACS buffer and counted using trypan blue exclusion method. Cells were then analyzed by flow cytometry as described previously (Flow Cytometry Analysis).

### Characterization of dendrimer size distribution and ζ-potential and dendrimer-serum protein interaction

The physicochemical properties (size distribution and ζ-potential) of amine (NH_2_), hydroxyl (OH), and succinamic acid (SA) dendrimers were characterized using Litesizer 500 (Anton Paar Instruments) at 25°C. To measure the hydrodynamic radius based on dynamic light scattering (DLS), all dendrimers with different terminal groups were dissolved in 1×PBS buffer (pH = 7.4) at 1mg/mL. The dendrimer solutions were filtered through a 0.22µm 13mm polyether sulfone (PES) syringe filter (Cytiva) before the size measurement within a 1 mL cuvette (Sarstedt). To measure the ζ-potential, dendrimers were diluted in 1×PBS buffer (pH = 7.4) at 0.3mg/mL. The dendrimer solutions were also filtered before the ζ-potential measurement within an omega cuvette (Anton Paar). To assess the interactions between dendrimers and serum proteins, dendrimers were incubated with mouse serum from C57BL/6 mice (in-house generated) for 30min at 37°C at a concentration of 0.86 mg/mL to allow the formation of the dendrimer-serum protein complex. The dendrimer-serum protein or the serum protein solutions were then diluted in 1×PBS buffer (pH = 7.4) to reach a final concentration of 0.3mg/mL before assessment.

### Mice and *in vivo* tumor models

Wildtype C57BL/6 and transgenic CCR2^RFP/WT^/CX3CR1^GFP/WT^ C57BL/6 mice were bred in-house at the UF animal facility. CCR2 ^RFP/WT^/CX3CR1^GFP/WT^ were generated by cross-breeding CCR2 deficient mice (CCR2 ^RFP/RFP^[B6.129(Cg)-CCR2^tm2.1lfv^/J]) and CX3CR1 deficient mice (CX3CR1^GFP/GFP^[B6.129P-CX3CR1^tm1Litt^/J]). All procedures involving animal housing, care, and surgical procedures were following the guidelines of the University of Florida Institutional Animal Care and Use Committee.

To generate an orthotopic model of GL261 and KR158 murine gliomas, mice were anesthetized by controlled isoflurane inhalation, and their heads were shaved before intravenous analgesia administration. Surgical sites were prepared using 2-3mm incisions at the midline of the skull. Stereotactic injection of 2µL at 1µL/min (5.0 x 10^4^) cells suspended in methylcellulose was performed at 2mm lateral from the bregma using a Hamilton syringe autonomously controlled by a micro-fluidic injection apparatus (Stoelting). Post-injection the dermal incision was closed via suture and bone wax application. Animals were placed on a cage warmer for post-surgical monitoring. For in vivo studies of dendrimer uptake and distribution, Mice received tail vein injections of SA and OH dendrimers (50mg/kg) and NH_2_ dendrimers (10mg/kg). Cy5-labeled dendrimers were suspended in 100-200µL saline and filtered with 0.22µm 13mm polyether sulfone (PES) filters.

### Flow cytometry sample preparation and analysis

Cells obtained from brain tumor, bone marrow (femur), spleen, and blood were analyzed by flow cytometry. Mouse blood was collected from the chest cavity post right atrium lancing using a 1mL syringe coated with 0.5M EDTA (Invitrogen). Approximately 200µL of blood was transferred to a 1.5mL microcentrifuge (Fisher Scientific) tube containing 100µL 0.5M EDTA (Invitrogen). Whole blood was centrifuged at 21°C, 380 x g for 5 minutes and the plasma was collected and stored at -80°C in 1.5mL microcentrifuge tubes. Before the collection of other organs, systemic perfusion was performed by needle insertion into the left ventricle and flushed with 20mL 1× PBS (Gibco) using a 10mL syringe (BD) and 25G butterfly infusion set (Exel). Brains were removed by sagittal and coronal partitioning of the skull using surgical scissors and transferred to a microscopy slide for tumor excision. To generate a single-cell suspension for analysis, tumor tissue was minced using a regular single-edge razor blade until a viscous suspension of cells was generated. Cells were then transferred to a 50mL conical (Falcon) filled with Accumax dissociation buffer (Innovative Cell Technologies) and incubated in a 37°C water bath for 5 minutes. Suspensions were then oscillated through a 1mL single-channel pipet tip for 40 cycles and strained through a 40µm strainer into a 50mL conical, followed by dilution with 5mL FACS buffer (2 or 10% FBS, 1× HBSS or 1X PBS). Cells were collected by centrifugation at 380 x g for 5 minutes at 4°C, followed by resuspension in 70% v/v Percoll Solution (GE) (70% Percoll, 1% 1× PBS in RPMI-1640). Using a 5mL syringe and 3-inch 18G needle, the 70% Percoll suspension of tumor cells was injected below a layer of 37% v/v Percoll (4mL, 37% Percoll, and 1% 1× PBS in Phenol-free RPMI-1640) (Gibco) in a 15 mL conical. Samples were subsequently centrifuged at 500 x g for 30 minutes at 21°C (level 1 acceleration, level 0 deacceleration). The resulting tumor cell interface between Percoll layers was removed (1mL) by a single channel pipet and transferred to a 1.5mL microcentrifuge tube. Cells were centrifuged at 500 ×g for 5 minutes at 4°C and washed and resuspended with ice-cold FACS Buffer (2 or 10% FBS in 1× HBSS (Gibco) or 1X PBS). Femurs were harvested and ends clipped with dissecting scissors after connective tissues were removed. The isolated femurs were placed in 0.5mL microcentrifuge tubes with an 18G needle pierced bottom, cap removed, and tube nested within a secondary 1.5 microcentrifuge tube containing 100µL ice cold ACK lysis buffer (Gibco). Microcentrifuge tubes with femurs were centrifuged at 5,700 RPM for 20 seconds at 21°C to capture bone marrow. Spleens were excised and transferred to a petri dish on ice for mincing using a regular single-edge razor blade (Personna) after injection with 1mL of ice-cold 1× HBSS (Gibco) or 1X PBS using a 3-inch 18G needle (Air-Tite) and 5mL syringe (BD). Dispersed tissues were aspirated into a 5mL syringe via a 3-inch 18G needle and transferred to a 15mL conical. 5mL of ice-cold 1× HBSS or 1X PBS was added, and cells were mechanically dissociated by the oscillation of the volume through the syringe and needle for 20 cycles. The resulting splenocyte suspension was centrifuged at 380 x g for 5 minutes at 4°C. 1mL ice cold ACK Lysis Buffer (Gibco) was added to bone marrow cells, leukocytes, and splenocyte to resuspend post centrifugation for 5 minutes and subsequently diluted with 5mL ice-cold FACS Buffer (2 or 10% FBS in 1× HBSS (Gibco) or 1X PBS) then strained through a 40µm cell strainer (Fisherbrand). Cells from each tissue were isolated by centrifugation at 380 x g for 5 minutes at 4°C. To remove all visibly present red blood cells, leukocytes were repeatedly cycled, up to four additional times, through ACK lysis buffer (Gibco) and FACS buffer wash as previously described. Viability was manually determined by cell count using a standard trypan blue (Corning) exclusion method.

Single-cell suspensions were prepared as described in the above sections. Samples were stained with viability dye (Violet, Invitrogen) in 1× PBS pH 7.4 (Gibco) at RT protected from light for 15 minutes. Cells were resuspended and washed with FACS buffer (2 or 10% FBS, 1× HBSS or 1X PBS) and stored on ice until analysis. Samples were analyzed via a single flow cytometry tube on a Sony Spectral Analyzer (SP6800). A multi-color reference control panel consisting of transgenic single color CCR2^RFP/WT^ and CX3CR1^GFP/WT^ bone marrow cells, viability dye–violet (Thermo Scientific), and Cy5-positive wildtype C57BL/6 bone marrow cell suspensions was utilized to unmix panels as appropriate. Raw data was subsequently analyzed and graphically illustrated using FlowJo software (BD Biosciences).

### *In vitro* dendrimer uptake study

Transgenic CCR2^RFP/WT^/CX3CR1^GFP/WT^ or WT C57BL/6 bone marrow cells were derived into M-MDSCs *ex vivo* as described herein (CCR2^RFP+^/CX3CR1^GFP+^). Cells were washed with FACS buffer (10% FBS, 1× HBSS) and resuspended in serum-free 1× HBSS (Gibco). Viability and concentration were determined by trypan blue exclusion. Cells were adjusted to a concentration of ∼1 × 10^6^/mL in 1.5mL microcentrifuge tubes (Fisher Scientific) and centrifuged at 500 x g for 5 minutes at 4°C. Cells were resuspended in 400µL of dissolved dendrimer solution at a concentration range from 1-100µg/mL in 1× HBSS or 1× PBS at room temperature and incubated for 30 minutes protected from light. Samples were then washed in 1× HBSS, stained for viability, and resuspended with FACS buffer in three technical repeats and analyzed via Spectral Flow Cytometry as described herein. To determine how dendrimer-associated serum proteins affect their interaction with M-MDSCs *in vitro*, dendrimer stock solutions were prepared by fully solubilizing dendrimers in 1× PBS pH 7.4 (Gibco), followed with filtration through a 13mm 0.22µm PES syringe filter (Cytiva). Solutions were diluted at room temperature to concertation of 0.86 mg/mL in either competent or heat-inactivated (60°C for 30 minutes) sex pooled, complement preserved, C57BL/6 murine serum (Charles River) in 1.5mL microcentrifuge tubes. Dendrimer-serum stock solutions were then incubated at 37°C for 30 minutes and brought to room temperature before co-incubation with cells at escalating doses (5-100µ/mL) in three biological repeats. The dendrimer uptake as indicated by Median Fluorescence Intensity (MFI) was subjected to flow cytometry analysis by gating out the CCR2^RFP+^/CX3CR1^GFP+^ (M-MDSC) or WT equivalent population.

### Immunofluorescence study

To determine the biodistribution and the cell uptake of dendrimers, Cy5-labeled dendrimers filtered through a 0.22μm sterile 13mm PES filter (Cytiva) were injected via tail vein into transgenic CCR2^RFP/WT^/CX3CR1^GFP/WT^ or wildtype C57BL/6 mice at tolerable doses (50 mg/kg for SA, OH dendrimers and 10mg/kg for NH_2_ dendrimers). Euthanized animals were systemically perfused with 20mL 1× PBS (Gibco) and 20mL 4% w/v paraformaldehyde (PFA) buffered solution. (Thermo Scientific) using a 50ml syringe and 25G butterfly needle infusion set. Brain (tumors), spleens, and femurs were excised, and connective tissues were removed prior to transfer to 5 mL of 4 % w/v PFA at 4°C for 1 hour (brains and spleens) or 72 hours (femurs) at 2–8°C. Brains (tumors) and spleens were then transferred to a 30% w/v sucrose (Fisher Scientific) in water (Corning) solution for ζ 24 hours in 15mL conical tubes stored at 2–8°C. Femurs were subsequently transferred to 5 mL of decalcification solution (20% EDTA, 10N NaOH, pH 7.4) for 4 days at 2-8°C and then a 30% w/v sucrose solution for 24 hours at 2-8°C. All tissues were embedded in optimal cutting temperature compound (Fisher Scientific) and cryo-sectioned at 10μm or 30μm thick sections at -25°C. Sections were prepared by addition to microscopy slides (Fisher Scientific), washed for 3 repetitive cycles with cold 1× Dulbecco’s Phosphate Buffered Saline (DPBS) in a staining dish. For vascular endothelial cell staining of brain tissues, an anti-mouse CD31-Spark YG 570 labeled mAb (BioLegend) was added to hydrophobic pen (Vector Laboratories) encircled sections at a concentration of 10 µg/mL. Slides were then mounted with Vectashield anti-fade mounting medium with DAPI stain (Vector Laboratories) and coverslip. Slides were sealed with CoverGrip sealant and subsequently stored at 2-8°C protected from light in the staining tray. Sections were analyzed at high magnification using an inverted Nikon A1R confocal microscope. Widefield fluorescent images were generated using a Keyence BZ-X800 or Nikon Ti-E for fluorescence microscopy. Widefield fluorescence microscopy images were processed using Nikon Elements software v5.21 and confocal fluorescence microscopy using Fiji v2.9.0.

### Quantification of dendrimer concentration in plasma

Blood was collected from euthanized CCR2^RFP/WT^/CX3CR1^GFP/WT^ mice bearing 3–4-week KR158 or GL261 gliomas at 24- or 72-hours post-dendrimer administration (tail vein) as previously described herein. Plasma samples were thawed from -80°C storage to room temperature and diluted 1:4 with 1× PBS (Gibco). Samples were then transferred to a 96-well clear bottom black plate (Thermo Scientific) and analyzed for absolute fluorescence intensity from Cy5 (635/675 (ex/em), integration: 400ms, read height: 3.0mm) using a Molecular Devices SpectraMax iD3 Multi-Mode Microplate Reader. Samples were plotted against a standard curve of Cy5 in murine serum and interpolated post-transgenic murine plasma background subtraction. The percentage of the injected dose was calculated by dilution factors × estimated dendrimer concentration × plasma volume (estimated to be 1.8 mL/mouse) / total injected dose. Samples were evaluated by 3 technical repeats.

### Data reporting and statistical analysis

Each *in vitro* assay was performed using a minimum of three technical repeats and three biological repeats. All data was processed and graphed using GraphPad Prism v10.1.1 displaying average, standard deviations, error, and statistical p-values by one or two-way ANOVA as appropriate per data set. Each in vivo assay was performed using a minimum of n=6 mice based on the median of a group comparison using a one- or two-way ANOVA between the calculated min and max degrees of freedom ((DF=k(n-1) n=sample size, k=number of groups). Representative immunohistochemistry n=1 (IHC) and microscopy tissue samples were displayed.

## Results and Discussion

### M-MDSCs represent a significant population in the glioma TME

To determine how dendrimers interact with M-MDSCs and other immune cells *in vivo*, we first characterized the profiles of M-MDSCs and other infiltrative immune cells in a GL261 mouse glioma model. This model well-recapitulates the histology of glioma and has been extensively used to test the therapeutic responses in the literatures^32^. GL261 gliomas were established in CCR2^RFP/WT^CX3CR1^GFP/WT^ transgenic mice, which allows the direct surveillance of the profile of infiltrative immune cells via the endogenously expressed red fluorescent protein (RFP) for chemokine receptor two (CCR2) and green fluorescent protein (GFP) for CX3C motif chemokine receptor 1 (CX3CR1)^20, 33^. When expressed jointly, these G-Protein Coupled Receptors (GPCRs) have been established as an equivalent biomarker for the M-MDSC cell subset, as defined by CD45^+^Ly6G^-^ Ly6C^+^CD11b^+^ populations^2^. These cells have been shown to suppress both CD4+ and CD8+ T cells in mouse glioma model^31^. At 2-3 weeks after the tumor initiation, we performed the flow cytometry analysis of the M-MDSC population from bone marrow, blood, and spleen. Our results showed that M-MDSCs accounted for 11.4±0.4% and 3.8±0.1% of total cells in the bone marrow and spleen of GL261 tumor-bearing mice (**Fig 1A and B**, orange). CCR2 and its cognate receptors mediated M-MDSCs egress from bone marrow into peripheral blood^31^, in which the M-MDSCs comprised 8.5±1.5% of the blood leukocytes (**Fig 1A and B**, orange); M-MDSCs infiltrated the glioma through peripheral blood, ultimately comprising 22.1±1.0% of the stromal cells in the GL261 tumor (**Fig 1C**, orange). In the TME, M-MDSCs were shown as the RFP and GFP co-localized cells, as indicated by the arrows in **Fig 1D**. The CCR2^RFP/WT^CX3CR1^GFP/WT^ transgenic mice also enabled us to profile other immune cell subsets in the glioma TME. Based on previously published data^20^, the CCR2^-^/CX3CR1^+^ subsets (16.7±3.0%, **Fig 1C**, dark green, abbreviated as microglia) were CD45^low^/MHC^+^/F4/80^+^/CD11c^-/^CD11b^medium^, likely representing the CNS tissue-resident microglia; the CCR2^+^/CX3CR1^-^ subsets (4.4±2.0%, **Fig 1C**, red, abbreviated as CCR2^+^) were CD45^+^/MHCII^+^/F4/80^-^/CD11c^-^/CD11b^low^, likely representing another myeloid cell that originated outside of the CNS; the CCR2^-^/CX3CR1^-^ subsets (26.4±4.9%, **Fig 1C**, grey, abbreviated as other cells) were a collection of tumor cells and other tumor stroma cells. Finally, the CCR2^-^/CX3CR1^int^ subsets accounted for 30.5±5.9% of tumor stromal cells (**Fig 1C**, light green, abbreviated as CXCR1^int^). Given these cells were a mixed population of CD45^low^ and CD45^high^ ^20^, it is possible that they infiltrated the glioma from outside of the brain.

**Figure 1.**
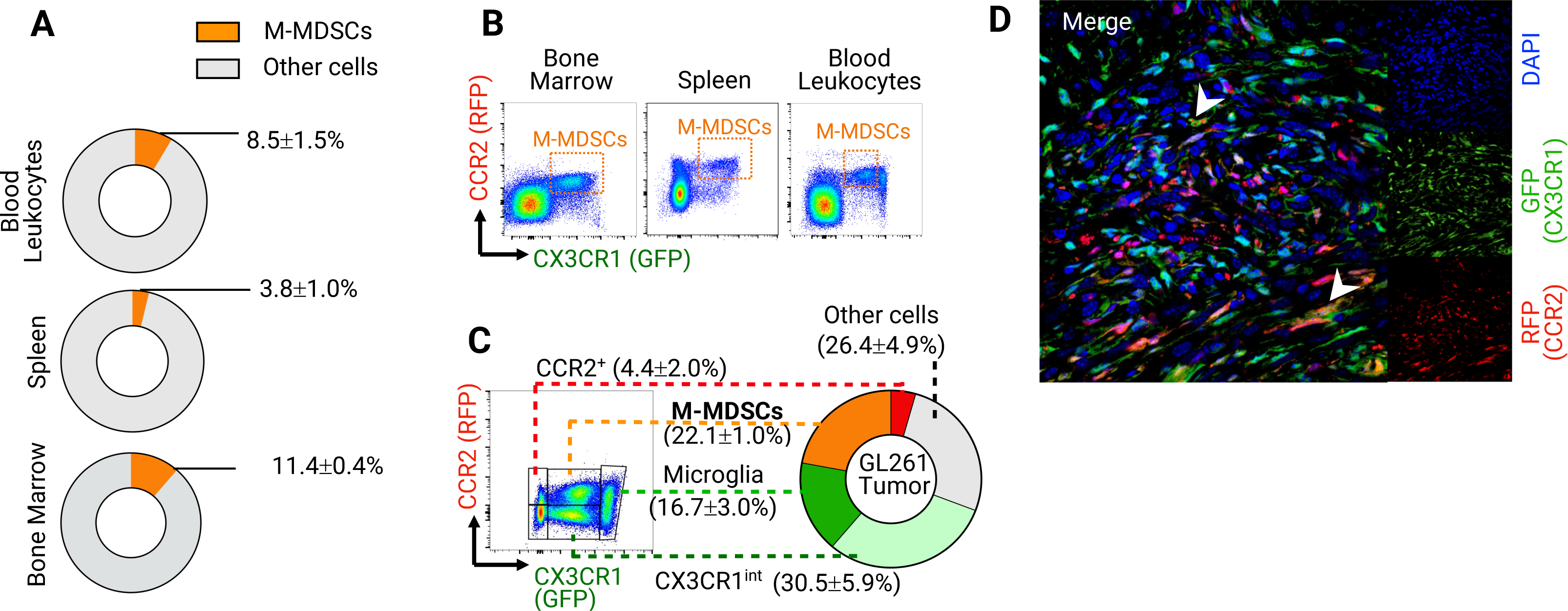
The CCR2^RFP/WT^CX3CR1^GFP/WT^ transgenic mice enable direct surveillance of M-MDSCs in mouse glioma model. **A)** in mice with established GL261 tumors, the pie graph shows the flow cytometry measurement of the average percentage of M-MDSCs reside in the reservoir tissues such as bone marrow, spleen, and peripheral blood leukocytes. At 3-4 weeks after tumor initiation, tissues from 6 tumor-bearing mice were analyzed. **B)** Gating strategy for the M-MDSCs cells in each tissue: the M-MDSCs are defined as the CCR2+/CX3CR1+ population, which is indicated in the orange box. **C)** Gating strategy and the average percentage of major cell subsets in the GL261 tumor stroma. CCR2+/CX3CR1+ cells (orange): M-MDSCs; CCR2-/CX3CR1+ cells (dark green): likely representing the CNS tissue-resident microglia; CCR2-/CX3CR1^meidum^ cells (light green): CX3CR1^int^, likely represent immune cells infiltrate brain tumor from external sources; CCR2+/CX3CR1- cells (red): CCR2+, likely represent other infiltrate myeloid cells originated outside of the CNS. CCR2-/CX3CR1- cells (grey): other cells, a collection of tumor cells and other tumor stroma cells. At 3-4 weeks after tumor initiation, tissues from 6 tumor-bearing mice were analyzed. **D)** confocal microscopy image of tumor (KR158). Arrow indicates the CCR2+/CX3CR1+ M-MDSCs. Red: RFP/CCR2; Green: GFP/CX3CR1; Blue: DAPI.

### Tumor M-MDSCs show a high capacity for dendrimer uptake

Previous studies have established that, in the presence of neuroinflammation/tumor lesions, systemically injected OH dendrimers can selectively localize in activated glial cells in a spectrum of central nervous system (CNS) disorders^28–30^. We evaluated OH dendrimer (Generation 6) as a model dendrimer to probe the dendrimer uptake capacity of different cell subsets within the glioma TME, the lymphoid organs (bone marrow, spleen), and the blood. Mice with established GL261 gliomas were injected systemically with OH dendrimers at 50mg/kg – a dose that has been well-tolerated *in vivo*^34^. To track the dendrimer–cell interaction, we fluorescently labeled the OH dendrimer with a minimal amount of Cy5 dye (∼5% by wt%)^29^. At 24 hours after injection, different cell subsets within the stroma of the GL261 tumor were isolated and subsequently subjected to flow cytometry analyses to determine the Cy5 Median Fluorescence Intensity (MFI) within each cell subset, which indicates the amount of dendrimer being endocytosed by the cells. Remarkably, tumor M-MDSCs and microglia showed a capacity for high dendrimer uptake (**Fig 2A**). Specifically, the MFI of tumor M-MDSCs =1061±535, which was significantly higher than CX3CR1^int^, CCR2+, and other cells (**Fig 2B**). This indicated that tumor M-MDSCs have a higher capacity for endocytosing OH dendrimers than other cell subsets within the GL261 tumor. We next evaluated the composition of all dendrimer-positive cells within the GL261 tumor by gating out the dendrimer-positive populations from the whole tumor stroma cells. The composition of the dendrimer-positive populations was then analyzed based on the CCR2 (RFP) and CX3CR1 (GFP) expression (**Fig 2C**). Our results showed that the majority of the dendrimer-positive cells were mostly distributed within 4 cellular compartments, i.e., M-MDSC (19.7±6.7%), microglia (25.1±3.4%), CX3CR1^int^ (28.6±6.9%), and other cells (24.0±6.2%) (**Fig 2D**). The CCR2+ compartment only accounted for 2.7±0.7% of dendrimer-positive cells (**Fig 2D**), potentially due to their small numbers within the tumor stroma (∼5%, **Fig 1C**).

**Figure 2.**
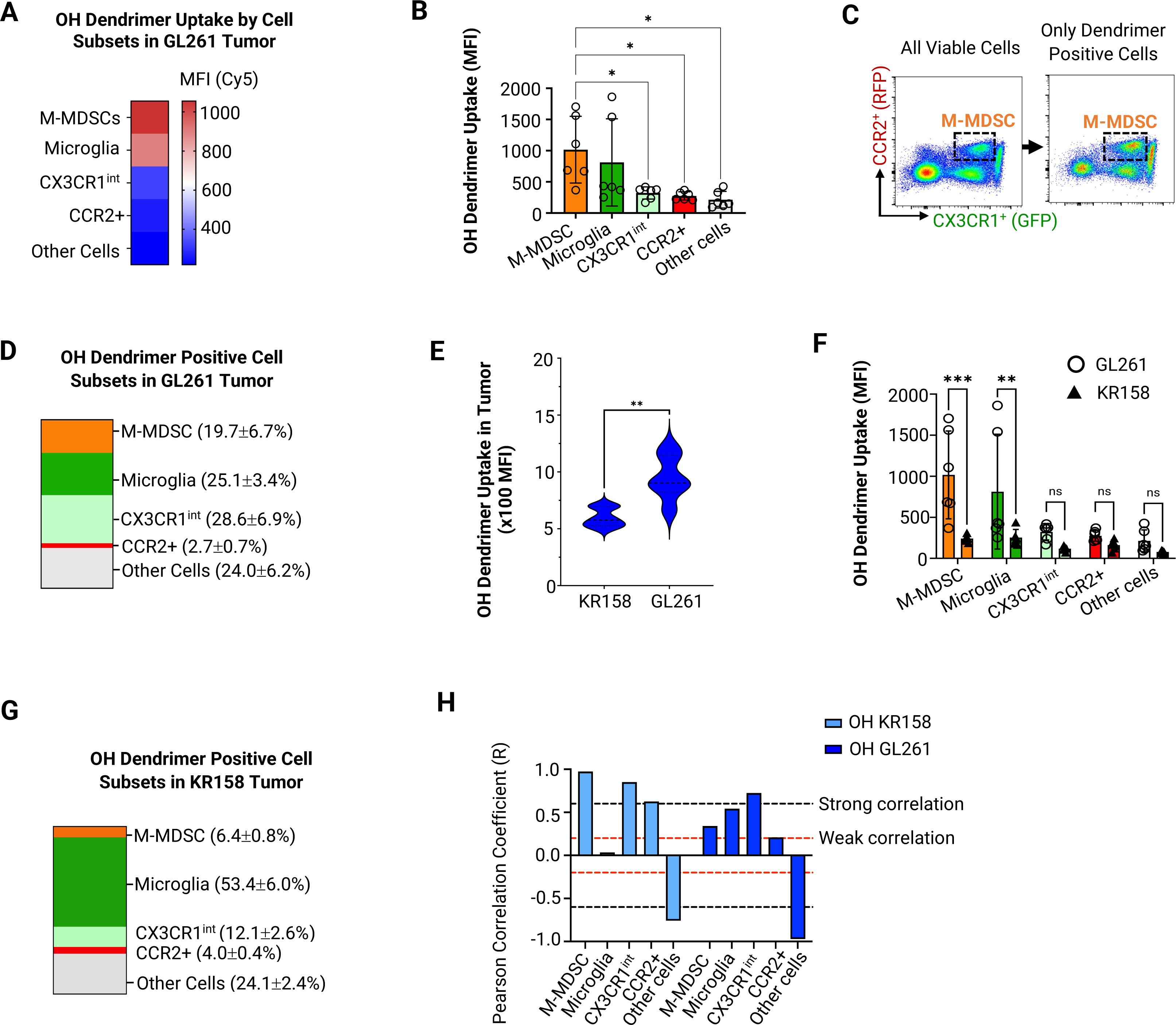
Tumor M-MDSCs efficiently endocytose dendrimers with high capacity. Cy5-labeled OH dendrimers (50mg/kg) were systemically injected into GL261 and KR158 tumor bearing transgenic mice (CCR2^WT/RFP^CX3CR1^WT/GFP^) at 3-4 weeks post-implantation; at 24 hours post injection, OH dendrimers uptake in myeloid subsets (indicated by Cy5 Median Fluorescence Intensity (MFI)) and the composition of dendrimer-positive cells (indicated by percentage of dendrimer-positive subset) within the tumor were analyzed through flow cytometry. **A)** The capacity of each cell subsets to uptake OH dendrimers within the GL261 gliomas at 24 hours post-injection. This is indicated by the MFI, which representatively measures the median number of dendrimers deposited per single cell. **B)** The statistical analysis of A). **C)** Representative density plot of GL261 tumor stromal cells (left panel) and in the same tumor, the density plot of OH dendrimer-positive cells (right panel). M-MDSCs are shown in the box. **D)** Statistical analysis of the density plot of dendrimer-positive cells in C) (right panel), which shows the composition of dendrimer-positive cells in each cellular compart within the GL261 tumor as a percent of a whole (n=6). **E)** Comparison of the mean, upper and lower quartiles of OH dendrimer deposition in tumor as measured by MFI between GL261 and KR158 gliomas. **F)** Comparison of OH dendrimer uptake capacity (MFI) in each cell subsets between GL261 and KR158 tumors. **G)** Composition of dendrimer-positive cells in each cellular compart within the KR158 tumor as a percent of a whole (n=6). **H)** the correlation analysis between the abundance of cell subset (indicated by the percentage of cell subset within all tumor stroma cells) and the dendrimer deposition within the tumor (indicated by Cy5 MFI of all tumor cells) for both GL261 and KR158 tumors. The Pearson correlation coefficient (R) is based on 95% confidence interval. Weak correlation, R>0.2 or R<-0.2; strong correlation, R>0.6 or R<-0.6. For all *in vivo* experiment, data were generated based on 6 mice of both male and female sexes, *p<0.05, **p<0.01, ***p<0.001, ns=not statistically significant

While the GL261 glioma model recapitulates the histology of glioma^32^, it is well-established that the GL261 glioma model, unlike human glioblastoma, is immunogenic^32, 35, 36^. Specifically, GL261 has high MHC-I expression and a high tumor mutational load^35^ and responds well to checkpoint inhibitors^36^. We next sought to characterize the dendrimer interactions with M-MDSC and other immune infiltrative cells in an KR158 model with lower populations of infiltrative M-MDSC and are resistant to checkpoint inhibitors^37, 38^. Interestingly, when comparing the overall Cy5 MFI of all tumor stromal cells, the GL261 tumor showed 1.6-fold higher MFI (mean=946.8) than the KR158 model (mean=598.2), indicating a higher dendrimer deposition in the GL261 tumors as compared to the KR158 tumors (**Fig 2E**). The different dendrimer deposition between the two glioma models was also reflected at the cellular level. The cell subsets within the GL261 tumor showed approximately 2–3-fold higher dendrimer uptake than the KR158 tumor (**Fig 2F**). Surprisingly, when comparing the cellular composition of dendrimer-positive populations, around 53.4±6.0% of dendrimer-positive cells in the KR158 tumor were located in the microglia compartment, while only 6.4±0.8% and 12.1±2.6% of dendrimer-positive cells were located in the M-MDSC and CX3CR1^int^ compartments respectively (**Fig 2G**). Although the KR158 tumor had lower dendrimer deposition and different compositions of dendrimer-positive cells compared to the GL261 tumor, the M-MDSCs and microglia in both tumor models showed higher dendrimer uptake than other cell subsets (**Fig 2F, Fig S1A-C**). In summary, our data based on two different glioma models indicated that the monocytic myeloid cells largely contributed to the tumor depositions of OH dendrimers. We next sought to evaluate whether the percentage of each cell subset within the tumor correlated with the amount of dendrimer deposition in both tumor models. Analyses of the Pearson correlation coefficients showed that the percentage of tumor-infiltrative cells (M-MDSCs, Microglia, CX3CR1^int^, and CCR2+) generally had positive correlations with the amount of OH dendrimer depositions in both tumor models (**Fig 2H)**. Specifically, M-MDSCs and CX3CR1^int^ population showed better correlation (R> 0.2 or R>0.6) than other cell subsets in both tumor models. However, the percentage of CCR2^-^/CX3CR1^-^ subsets (other cells), which are a collection of tumor cells and other tumor stroma cells, showed a strong negative correlation with OH dendrimer deposition (R<0.6).

The selective uptake of OH dendrimer by tumor-associated microglia/macrophages has been reported in previous studies^28, 29^. However, ontogeny differences between CNS-resident microglia and bone marrow-derived macrophages suggests distinct functions in brain cancer and different responses to macrophage-targeting therapeutics^39, 40^, which highlights the importance of analyzing the cell-level biodistribution of nanotherapeutics. Both the M-MDSCs originated from bone marrow and the CNS-resident microglia showed a strong capacity for taking up OH dendrimers in our study. However, it is possible these cell subsets endocytose dendrimers through different mechanisms. For example, bone marrow-derived M-MDSCs and tumor-associated macrophages are more associated with phagocytosis and antigen-presentation. They are present at a higher density in the perivascular niche than microglia, which display signatures associated with synaptic pruning^41, 42^. We also noticed the differential dendrimer deposition between the GL261 and KR158 tumors. This difference might be associated with the different immunogenicity of GL261 and KR158 tumors^32^. The immunogenic GL261 tumor has a ‘hotter’ tumor milieu with more infiltrative immune cells than KR158 tumors^32^. Since ours and other studies showed infiltrative myeloid cells within the tumor often contribute to the tumor-accumulation of nanoparticles^43, 44^, it is possible that the higher OH dendrimer deposition in the GL261 tumor was associated with the higher amount of infiltrative immune cells in this model. The KR158 tumor showed a mushroom-like crown on top of the brain, while the GL261 histology was more representative of the human glioma (**Fig S2**). As the tumor pathophysiology can also significantly impact nanoparticle deposition^45^, it remains to be determined whether the histological difference could also contribute to the differential dendrimer uptake.

### The trafficking kinetics of M-MDSC contribute to the dendrimer accumulation in the tumor

In the presence of tumor lesions, the production of M-MDSCs is accelerated in the bone marrow, from which these cells are directly recruited to the brain tumor through peripheral blood or indirectly from the spleen, which serves as the temporary reservoir of M-MDSCs^20, 31, 33^ (**Fig 3A**). To determine whether the trafficking kinetics of M-MDSCs affected the deposition of dendrimer in the brain tumor, we first quantified the cellular uptake of OH dendrimers (MFI) by M-MDSCs located in bone marrow, spleen, peripheral blood, and tumor (GL261) at 24 hours after dendrimer injection (50mg/kg). Dendrimer uptake was observed in the M-MDSCs from all tissues analyzed (**Fig 3B**). In the femur bone, OH dendrimers were mostly distributed in the red marrow, where hematopoiesis led to the product of leukocytes (**Fig S3A**). In the spleen, OH dendrimers were mostly distributed in the red pulp (**Fig S3B**), where MDSCs are located^46^. The blood M-MDSCs showed the highest dendrimer uptake (MFI=1371±494), probably because blood M-MDSCs can directly access the dendrimers in the circulation without the limitation of any tissue barriers. We further analyzed the dendrimer uptake in different blood leukocytes (identified through the FSC- and SCC-based scattered plots). The dendrimer uptake was highest in granulocytes (MFI=1112±232), followed by monocytes (MFI=231.7±48) and lymphocytes (MFI=172.8±56) (**Fig 3C** and **Fig S4A**). Approximately 95% of granulocytes showed dendrimer uptake, compared to ∼40% for monocytes and ∼30% for lymphocytes (**Fig S4B)**. Since M-MDSCs are constantly being recruited to the tumor in large amounts during tumor development, we hypothesize that blood M-MDSCs may carry endocytosed dendrimer to the tumor while they infiltrate the tumor stroma. To test this hypothesis, we analyzed the change of the dendrimer-positive M-MDSCs percentage between the 24- and 72-hour window in two cohorts of mice. In bone marrow, we observed a 50% decrease in the percentage of dendrimer-positive M-MDSCs within the 48-hour window (**Fig 3D**). This decrease in dendrimer-positive M-MDSCs could be caused by two factors. First, the emergency myelopoiesis in cancer leads to the accelerated generation of new M-MDSCs, which could dilute the dendrimer-positive populations; Second, the initial dendrimer-positive M-MDSCs could egress from the bone marrow, further diluting the percentage of dendrimer-positive M-MDSC in the bone marrow. In the GL261 tumor, we observed a 90% increase in the percentage of dendrimer-positive M-MDSCs (**Fig 3D**). This significant increase in dendrimer-positive M-MDSCs may have resulted from the recruitment of external dendrimer-positive M-MDSCs to the tumor milieu during the 48-hour window. It is less likely that tumor M-MDSCs could uptake more dendrimers within the 48–72 hours window, as further analysis of serum concentration showed that the amount of OH dendrimers decreased from 12.5±8.5% (total injected dose) at 24 hours to only 4.6±3.7% at 72 hours (**Fig 3E**). There was no significant change in dendrimer-positive populations in the blood and spleen M-MDSCs.

**Figure 3.**
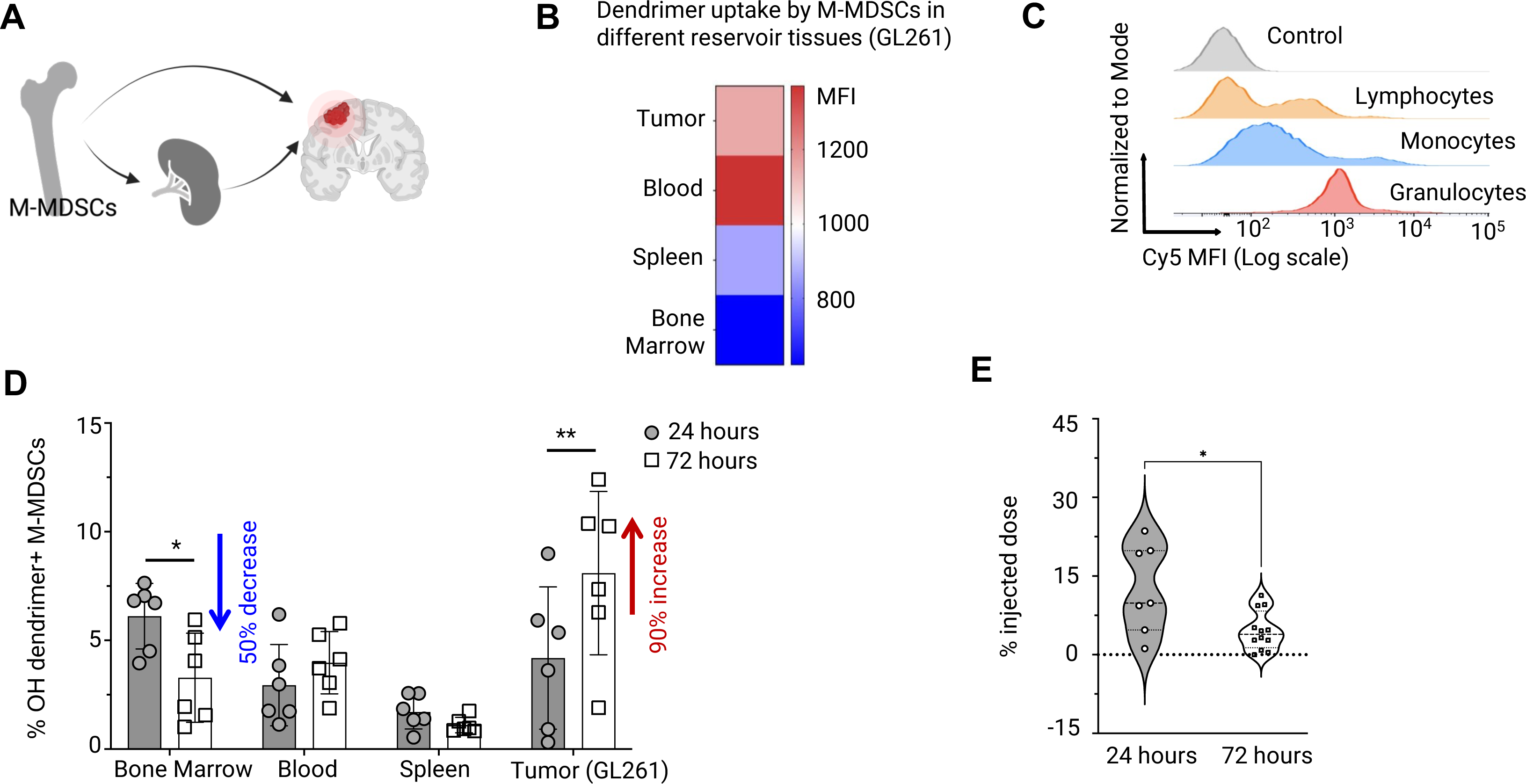
The trafficking kinetics of M-MDSC contributes to the dendrimer accumulation in the tumor. **A)** Graphical illustration shows the trafficking kinetics of M-MDSCs. M-MDSCs are recruited to the brain tumor from bone marrow (hematopoiesis organ) or spleen (temporary reservoir) through systemic circulation (blood). **B)** Heatmap shows the OH dendrimer uptake capacity (indicated by Cy5 MFI) for M-MDSCs in bone marrow, spleen, blood, and tumor in GL261 glioma bearing mice (average MFI from 6 mice). **C)** Histogram shows the overall OH dendrimer uptake capacity (indicated by Cy5 MFI) for white blood cells isolated from the peripheral blood of GL261 tumor-bearing mice. Control cells (no dendrimer injection). Cell subsets (lymphocytes, monocytes, granulocytes) were gated based on the scattered plot FSC vs. SSC. **D)** Comparison of the percentage of dendrimer-positive M-MDSCs between 24 hours vs. 72 hours after dendrimer injection. Data was obtained from GL261 tumor-bearing mice. **E)** Quantification of OH dendrimer concentrations in the plasma of GL261 tumor-bearing C57BL/6 mice at 24 hours (n=7) and 72 hours (n=12) post-dendrimer injection. For all other experiments in this figure, flow cytometry analyses were based on 6 GL261 tumor-bearing mice that received systemic injection of 50 mg/kg OH dendrimers. *p<0.05, **p<0.01.

For decades, the Enhanced Permeability and Retention (EPR) effect has been used as the guiding principle for designing tumor-targeting nanomedicine^2^. Previous studies based on this theory showed that the accumulation of OH dendrimers in glioma was largely attributed to their neutral surface and ultra-small sizes (sub-10nm), which allowed them to efficiently cross the impaired blood-brain tumor barriers (BBTB) taken up by myeloid cells within the tumor through EPR effect and^28, 29^. However, it was not clear whether other complementary mechanism(s) could also contribute to the dendrimer accumulation in the tumor. Historic studies demonstrated myeloid cells can modulate the pharmacokinetics, biodistribution, and efficacy of nanotherapeutics^12, 13^. Recent studies argued that circulating myeloid cells, such as inflammation-associated monocytes and granulocytes can actively transport nanoparticles from the blood to the tissue when they infiltrate the inflamed tissue^14, 43, 47^. Our study confirmed this alternative mechanism in a mouse model of glioma, by showing that highly tumor-infiltrative M-MDSCs can contribute to the tumor accumulation of OH dendrimers. This phenomenon can be leveraged to design M-MDSC-targeting therapeutics for enhanced tumor delivery. Future studies in this direction could benefit from a quantitative study of the tumor accumulation of adoptively transferred dendrimer-positive M-MDSCs.

### Dendrimer surface functionality affects their interactions with M-MDSC *in vivo*

Dendrimer surface functionality can significantly affect their *in vivo* behaviors, such as absorption, distribution, metabolism, elimination (ADME), and toxicity^30, 48–50^. We next asked how the cell-level distribution of systemically injected dendrimers can be affected by their surface chemistry. Herein, we selected G6 PAMAM dendrimers with succinamic acid (SA), hydroxyl (OH), and amine (NH_2_) terminal groups. All three dendrimers showed approximately ∼6 nm diameter (number average mean, **Fig S5A**). When measured in 1×PBS, NH_2_, OH, and SA dendrimers showed cationic (ζ=32.2±0.5mV), neutral (ζ=5.0±0.2mV), anionic (ζ=-22.4±0.6mV) surface charges (**Fig S5B)**. To trace dendrimers *in vivo*, all three dendrimers were stably labeled with a minimal amount of Cy5 (5% by wt.%) through previously established conjugation chemistry^30^ (**Fig S5C)**. We started by systemically injecting all three dendrimers (50mg/kg) into transgenic mice with established KR158 tumors or age-matched healthy controls. At the dose of 50mg/kg, NH_2_ dendrimers induced significant toxicities that ultimately led to animal death. The high *in vivo* toxicity of systemically administrated NH_2_ dendrimers has been reported previously^51^. Our *in vitro* toxicity study based on bone marrow-derived M-MDSCs also showed when the dendrimer dose is ≥ 20µg/mL, NH_2_ dendrimers showed significantly higher toxicity than OH and SA dendrimers (**Fig S6A)**. Therefore, we lowered the dose of NH_2_ dendrimers to 10mg/kg for the following *in vivo* studies.

### NH_2_ dendrimers cannot efficiently access M-MDSCs, but can be readily taken up by M-MDSCs

Nanoparticles need to efficiently cross tissue barriers (e.g. the blood vessels and tissue extracellular matrix) before successfully accessing the cells located in the tissue stroma. We determined how dendrimer surface functionality affects their abilities to selectively be endocytosed by M-MDSCs by measuring the percentage of dendrimer-positive M-MDSCs in tissue (% dendrimer+ M-MDSCs). The bone marrow and the tumor are the origin and the destination of M-MDSC recruitment, therefore were selected as the tissues of interest in this study. We found that the percent of dendrimer+ M-MDSCs was significantly lower for NH_2_ than other dendrimers in the bone marrow of both KR158 tumor-bearing mice and healthy control (**Fig 4A)**. In the KR158 tumor stroma, NH_2_ dendrimers also targeted less M-MDSCs (12.3±5.3%) than SA (28±12.7%) and OH dendrimers (19.0±6.3%) (**Fig 4B)**. The lower cell-targeting of NH_2_ dendrimers was likely due to their lack of ability to cross tissue barriers^30^. Confocal imaging of KR158 tumor sections showed that NH_2_ dendrimers were mostly co-localized with the endothelial cells along the blood vessels (**Fig S6B and S7A**), indicating the NH_2_ dendrimers with strong cationic surface charge, were not able to cross the BBTB and other tissue barriers^30^. However, OH and SA dendrimers were able to efficiently cross the BBTB and distribute within the tumor stroma (**Fig S7 D, E, F and G, H, I**). Interestingly, although NH_2_ dendrimers were not taken up by many M-MDSCs in the bone marrow, the M-MDSCs that endocytosed NH_2_ dendrimers showed similar quantities for tumor-bearing mice and higher quantities of intracellular dendrimers for healthy controls over OH and SA dendrimers (**Fig 4C**). Specifically, in the bone marrow of healthy mice, the NH_2_ dendrimer (MFI=1236±453) showed 3.2-fold and 2.7-fold higher M-MDSC-uptake than OH (MFI=384±15) and SA dendrimers (MFI=466±33) (**Fig 4D**). In the tumor M-MDSCs of KR158 tumor-bearing mice, NH_2_ dendrimers also showed slightly higher MFI (MFI=685±172) than OH (MFI=442±144) and SA dendrimers (MFI=595±103) (**Fig 4E**, not statistically significant). Given the dose of NH_2_ dendrimer is 5-fold lower (10mg/kg) than OH and SA dendrimers, NH_2_ dendrimers showed higher capacity for M-MDSC uptake.

**Figure 4.**
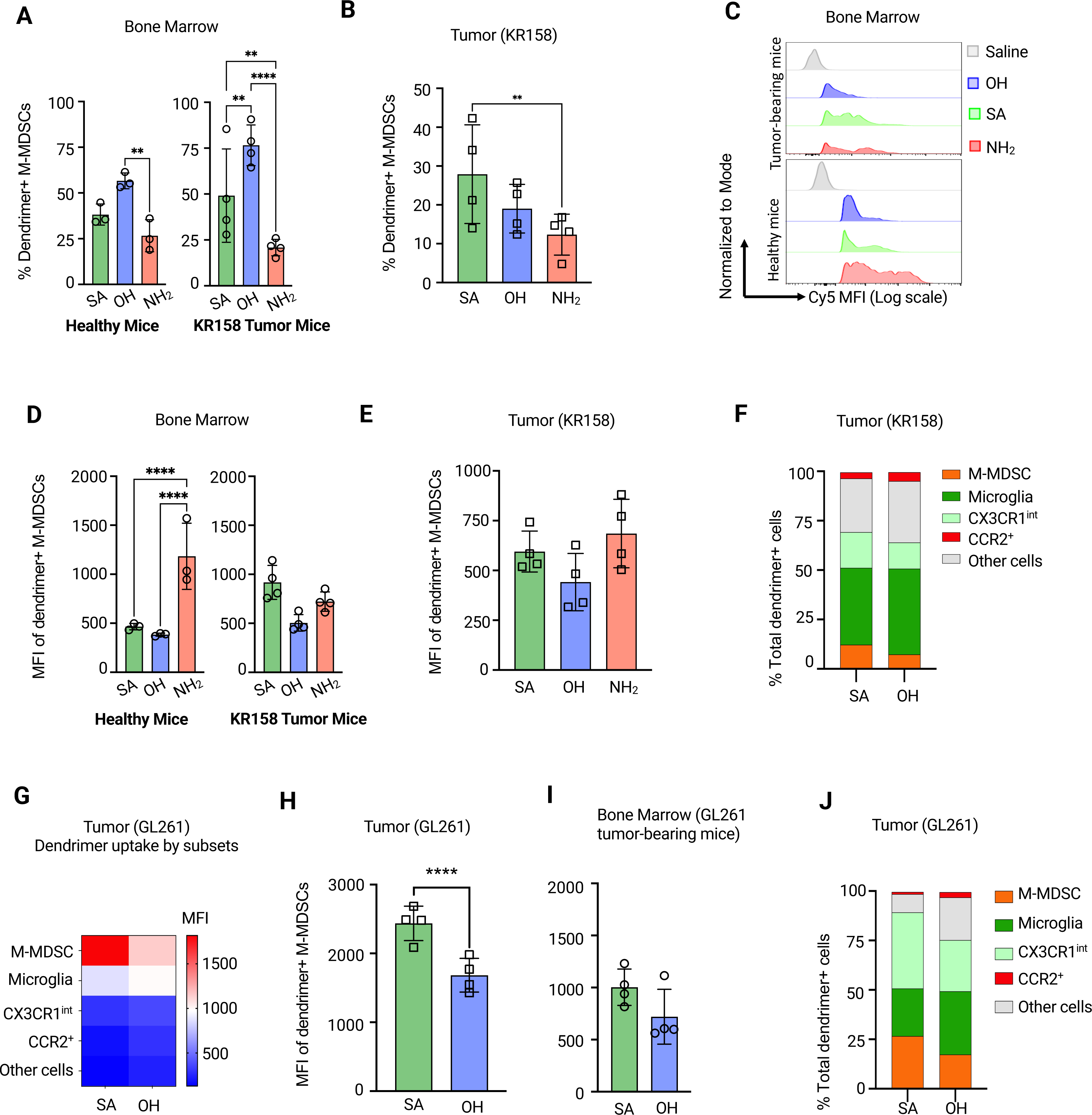
Dendrimer surface chemistry affects their interactions with M-MDSC *in vivo.* To determine how dendrimer surface chemistry affect their interaction with M-MDSCs in vivo, dendrimers with different terminal groups (i.e. Succinamic acid: SA, hydroxyl: OH, and amine: NH_2_) were systemically injected into healthy (n=3) or tumor bearing mice (n=4) at tolerable doses (50mg/kg for SA and OH, 10mg/kg for NH_2_). At 24 hours after dendrimer injection, the bone marrow and tumor were isolated for flow cytometry analysis. **A)** comparison of the percentage of dendrimer-positive M-MDSCs (% dendrimer+ M-MDSCs) in the bone marrow of healthy (left) and KR158 tumor-bearing mice (right) between SA, OH, and NH_2_ dendrimers. The % dendrimer+ M-MDSCs measure dendrimer’s ability to ‘target’ M-MDSCs in the tissue. **B)** comparison of the percentage of dendrimer-positive M-MDSCs in the KR158 tumor between SA, OH, and NH_2_ dendrimers. **C)** Representative histograms comparing the Cy5 MFI of dendrimer-positive cells for OH, SA, and NH_2_ dendrimers in the bone marrow of KR158 tumor-bearing mice (top) and healthy mice (bottom) respectively. **D, E)** Comparison of the M-MDSC’s capacity to endocytose SA, OH, and NH_2_ dendrimers (indicated by MFI). Plot shows M-MDSCs isolated from D) the bone marrow of healthy mice (left) and KR158 tumor bearing mice (right) and **E)** the KR158 tumors. **F)**. Comparison of SA and OH dendrimers for the composition of dendrimer-positive cells within the KR158 tumor. **G)** the heatmap shows the uptake capacity of SA and OH dendrimers (indicated by Cy5 MFI) by different cell subsets within the GL261 tumor. Data displayed is median. **H, I)** Statistical analysis comparing the SA and OH dendrimers in terms of their uptake capacity (indicated by Cy5 MFI of dendrimer-positive cells) by M-MDSCs within the tumor (H) and bone marrow (I) of the GL261 tumor bearing mice. **J)** Comparison of SA and OH dendrimers for the composition of dendrimer-positive cells within the GL261 tumor. For all statistical analyses, *p<0.05, **p<0.01, ****p<0.0001. ns: p value 20.05 was not shown.

### M-MDSCs take up SA dendrimers more readily than OH dendrimers

Although SA and OH dendrimers did not show significant differences in their overall tumor deposition **(Fig S8)**, they did display differential cellular uptake by M-MDSCs. Specifically, in the KR158 tumor, M-MDSCs took up more SA than OH dendrimers (**Fig 4D and E**) 24 hours after injection. When comparing the MFI of dendrimer-positive M-MDSCs in the bone marrow of tumor-bearing mice, the SA (MFI=918±173) was 82% higher than OH (MFI=504±85) (**Fig 4D**), while for tumor M-MDSCs, the SA (MFI=595±103) was 35% higher than OH (MFI=442±144). In mice that received SA dendrimers, the M-MDSCs compartment and the CX3CR1^int^ compartment (cells that are potentially derived from M-MDSCs) accounted for a higher fraction (30.3%) of the dendrimer-positive cells when compared to mice that received OH dendrimers (20.5%) (**Fig 4F**). We further validated the difference between SA and OH dendrimer uptake in the GL261 tumor model. Within the GL261 tumor stroma, M-MDSCs took up more SA and OH dendrimers than other cell subsets (**Fig 4G**). Tumor M-MDSCs took up 45% higher SA (MFI=2437±250) than OH dendrimers (MFI=1683±245) (**Fig 4H**), while bone marrow M-MDSCs took up 40% higher SA (MFI=1003±175) than OH dendrimers (MFI=720±264) (**Fig 4I**). Similar to the KR158 tumor, in the GL261 tumor of SA dendrimer-injected mice, the M-MDSCs compartment and the CX3CR1^int^ compartment accounted for a higher fraction (65.1%) of dendrimer-positive cells than OH dendrimer injected mice (43.2%) (**Fig 4J**). In summary, these results showed that SA dendrimers were more efficiently endocytosed by M-MDSCs than OH dendrimers.

### Dendrimer-associated serum proteins mediate the interactions between dendrimer and M-MDSCs

Given that M-MDSC showed different capacities for endocytosing NH_2_, OH, and SA dendrimers *in vivo*, we next sought to determine the mechanism behind the differential uptake by testing the dendrimer uptake in *ex vivo* generated M-MDSCs. Adapted from an established ex vivo M-MDSC culturing model^31^, we exposed bone marrow cells isolated from CCR2^RFP/WT^CX3CR1^GFP/WT^ transgenic mice to KR158 conditioned media for 5 days. Flow cytometry analysis showed that we can expand the population of CCR2^+^/CX3CR1^+^ cells in the bone marrow from less than 10% to approximately 59% of the total cells *ex vivo* (**Fig 5A and B**). These cells successfully recapitulate the immune suppressive features and the migration pattern of M-MDSCs in the tumor-bearing mice^31^. When these *ex vivo* generated M-MDSCs were exposed to NH_2_, OH, and SA dendrimers in a serum-free media at a non-toxic dose of 10µg/mL (**Fig S6A**), a differential uptake of dendrimers was observed. Specifically, the dendrimer uptake was the highest for NH_2_, followed by SA, and OH dendrimers (**Fig 5C**).

**Figure 5.**
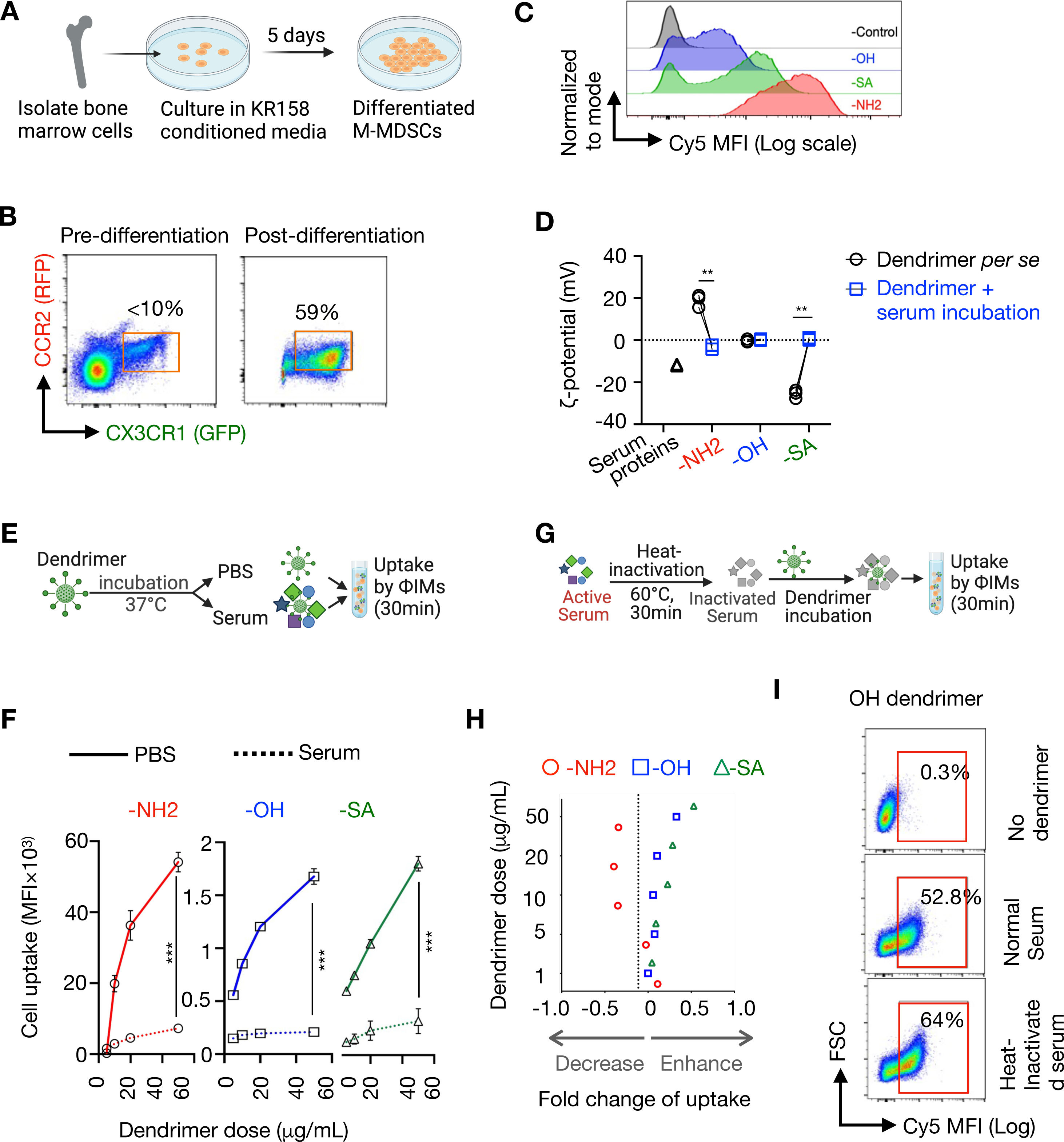
The serum proteins associated with dendrimers dictate the interaction between dendrimers and M-MDSCs. **A)** Schematic illustration shows the generation of M-MDSC from the bone marrow of CCR2^WT/RFP^CX3CR1^WT/GFP^ transgenic mice. To generate the M-MDSCs, bone marrow cells were isolated and were cultured in KR158 conditioned media for 5 days. **B)** flow cytometry analysis shows that exposure of bone marrow cells to KR158 conditioned media for 5 days enriched the CCR2+/CX3CR1+ cells (M-MDSCs) from less than 10% to approximately 59%. **C)** The histogram compares the uptake of OH, SA, and NH2 dendrimers (indicated by Cy5 MFI) by *ex vivo* generated M-MDSCs in the absence of mouse serum. To generate the plot, M-MDSCs were incubated with dendrimers at room temperature (RT) for 30 minutes. **D)** Comparison of ζ-potentials (mV) of serum proteins (black triangle) and NH2, OH, and SA dendrimers before (black circle) and after incubation with mouse serum (blue square). Dendrimers were incubated at 0.86 mg/mL in normal murine serum at 37°C for 30 minutes. ζ-potentials (mV) determined using DLS. The experiment was repeated for 3 times (n=3). **E)** Schematic illustration shows the experiment flow that determines the influence of serum proteins on the dendrimer uptake by M-MDSCs. **F)** the plot based on the experiment flow in (E), which shows dose-dependent dendrimer uptake (indicated by Cy5 MFI) by M-MDSCs after co-incubation with either PBS (solid line) or mouse serum (dotted line) for NH2 (red, circle), OH (blue, square), and SA (green, triangle) dendrimers. Each date point is an average of 3 independent experiment. **G)** Schematic illustration shows the experiment flow that determines how heat-inactivation of serum affect the uptake of NH2, OH, and SA dendrimers by M-MDSCs. **H)** the plot based on the experiment flow in (G), which shows the fold change of dendrimer uptake (x-axis) before and after heat-inactivation of mouse serum as a function of dendrimer dose (y-axis). NH2: red circle, OH: blue square, SA: green triangle. Negative fold change indicates heat inactivation of serum decreased dendrimer uptake. Positive fold change indicates heat inactivation of serum enhance dendrimer uptake. Each date point is an average of 3 independent experiment. **I)** Representative scattered plot shows heat inactivation of mouse serum enhanced the uptake of OH dendrimers by M-MDSCs. **p<0.01, ***p<0.005.

Although histologic studies have established that the surface charge of dendrimers can significantly affect their interaction with cells^49, 50, 52, 53^, it is now recognized that for systemically injected nanoparticles, it is the serum proteins that are associated with the nanoparticles that dictate nanoparticle interactions with cells^25, 54^. To determine how dendrimer-associated serum proteins affect their uptake by M-MDSCs, we first investigated how serum proteins interact with NH_2_, OH, and SA dendrimers by evaluating the change of ζ-potential after incubating dendrimers with mouse serum (0.86mg/mL). Interestingly, after 30min incubation under 37°C with mouse serum (ζ-potentials=-11.8±0.37mV), the original ζ-potentials of NH_2_ (32.2±0.5mV, cationic), OH (5.0±0.2mV, neutral), and SA (-22.4±0.6mV, anionic) all became neutral (NH2: -3.0±1.1mV; OH:0.3±0.6mV; SA: 0.5±0.5mV) (**Fig 5D**), indicating the surfaces of all three dendrimers were masked by the serum proteins. We then exposed all dendrimers either with or without serum incubation to bone marrow-derived M-MDSCs at an escalating dose of 5µg/mL to 50µg/mL (**Fig 5E**). Pre-incubating dendrimers with serum significantly decreased the uptake of all dendrimers in a dose-dependent manner, when compared to dendrimer uptake in non-serum containing PBS (**Fig 5F**), or HBSS media (**Fig S9C**) indicating that ‘native’ dendrimers and serum protein- ‘coated’ dendrimers interacted with M-MDSCs via different mechanisms. The uptake of NH_2_ dendrimers had a greater decrease after serum incubation than the OH and SA dendrimers (**Fig 5F, Fig S9A,B**), indicating the serum protein had a greater influence in mediating the interaction of M-MDSCs with NH_2_ dendrimers. Serum proteins can be classified into opsonin (enhance uptake) and dysopsonin (reduce uptake)^54^. To determine the classes of proteins that interreacted with dendrimers of different surface functionalities, we incubated dendrimers with either competent ‘active’ serum or ‘heat-inactivated’ mouse serum under 60°C for 30min (**Fig 5G**), for NH_2_ dendrimers, heat-inactivation of serum proteins reduced NH_2_ dendrimer uptake in a dose-depended manner for up to 60% (**Fig 5H, Fig S9 D-I**), indicating serum proteins associated with NH_2_ dendrimers actively mediated their uptake by M-MDSCs. However, for both OH and SA dendrimers, heat-inactivation of serum proteins increased their uptake by M-MDSCs up to 24% (OH) and 43% (SA) in a dose-dependent manner (**Fig 5H, Fig S9 D-I**), indicating a different class of serum proteins (potentially dysopsonins) might be associated with OH and SA dendrimers.

Historical studies of the ‘protein corona’ associated with nanoparticles are established on nanotherapeutics of 20 – 500nm size ranges with internally encapsulated payloads^26^, such as lipid NPs, polymers, and iron oxide NPs. Little is known, however, about NPs with ultra-small architectures, such as dendrimers in the 1 – 20nm size range. Compared to large NPs, dendrimers have sizes similar to proteins and may interact with serum proteins in different stoichiometries and configurations^54^; More importantly, dendrimers carry payloads on their surfaces. Therefore, the properties of the surface payload will affect how dendrimers interact with M-MDSCs. Here, we used NH_2_ and SA dendrimers to represent dendrimers carrying drug molecules of acidic and basic properties and compared them with OH dendrimers (control). We showed that the serum proteins associated with SA and OH dendrimers had a similar influence on their interaction with M-MDSCs, while certain serum proteins associated with NH_2_ dendrimers actively enhance their uptake by M-MDSCs, potentially through receptor-mediated endocytosis. In fact, a recent study showed that NH_2_ dendrimers efficiently interact with IgM and complement protein C3^55^, enhancing their phagocytosis. These *ex vivo* studies provided a basic mechanistic explanation for the differential *in vivo* uptake of NH_2_, OH, and SA dendrimers by M-MDSCs. However, to complete the mechanistic study, future research is needed to identify the specific proteins and their cognate receptors on M-MDSCs that mediate the uptake of dendrimers.

## Conclusion

M-MDSCs suppress the anti-tumor immune response locally at the TME and globally at the lymphoid organs. To address the systemic immune suppression, nanoparticles need to efficiently target these cells both locally and globally. Using the CCR2^RFP/WT^CX3CR1^GFP/WT^ transgenic mice, that enable direct surveillance of M-MDSCs, we showed that M-MDSCs can infiltrate glioma through peripheral blood in large amounts. Systemically injected hydroxyl dendrimers efficiently target M-MDSCs located in bone marrow, peripheral blood, spleen, and tumor. Within the tumor, M-MDSCs and microglia showed high capacity to endocytose hydroxyl dendrimers and these two cellular compartments accounted for more than half the amount of the hydroxyl dendrimer deposition in the tumor. In the GL261 glioma model, which has a high abundance of infiltrative immune cells, dendrimer showed greater tumor deposition and higher efficiency of M-MDSC-targeting than the KR158 glioma model. We further showed that the recruitment of M-MDSCs from bone marrow to tumor contributed to the tumor deposition of hydroxyl dendrimers. The surface functionality of dendrimers affects their ability to target M-MDSCs *in vivo*. Although amine dendrimers had the highest capacity of being endocytosed by M-MDSCs, they could not access these cells as efficiently as hydroxyl or succinamic acid dendrimers, potentially due to the lack of ability to cross tissue barriers. M-MDSCs took up succinamic acid dendrimers more efficiently than hydroxyl dendrimers. Finally, serum proteins can affect how dendrimers interact with M-MDSCs. The serum proteins associated with amine dendrimers significantly enhanced their uptake by M-MDSCs, while serum proteins associated with hydroxyl and succinamic acid dendrimers slightly reduced their uptake by M-MDSCs. Given that dendrimer-based drug conjugates carry drug payload on their surfaces, the results of this study indicated that the payload molecular properties could affect the *in vivo* fate, such as cell- and tissue-targeting of the final dendrimer-drug conjugates.

## Author Contributions

C.A.L. designed and performed the experiments, analyzed the experiment data, and helped draft the manuscript. G.P.T. designed and performed the experiments and analyzed the experiment data. C.S.S synthesized and characterized the fluorescently labeled dendrimers. A.S. induced gliomas in mice. K.A.R. and J.S.G. helped perform the *in vivo* experiments and analyzed the experiment data. J.K.H. developed and provided the mouse glioma models based on the transgenic mouse, helped conceive the study, and designed the experiments. F.Z. conceived the study, helped design the experiments and analyzed the experiment data, and wrote the manuscript.

## Acknowledgments

The authors are grateful to Ms. Wen Jiang for her help with the tissue sectioning and data validation and Ms. Tejashwin Lnu for her help with dendrimer physiochemical properties and dendrimer-serum interaction characterization. We also thank UF Pharmacology & Therapeutics, UF ICBR Cytometry Core (RRID:SCR_019119), and Electron Microscopy Core (RRID:SCR_019146) for providing the cryostat, flow cytometry, and confocal microscopy. This work is supported by the UF Research Opportunity Seed Fund and the American Brain Tumor Association Research Discovery Grant (DG2200050).

