## Supplementary Figures for "Systemically targeting monocytic MDSCs using dendrimers and their cell-level biodistribution kinetics"

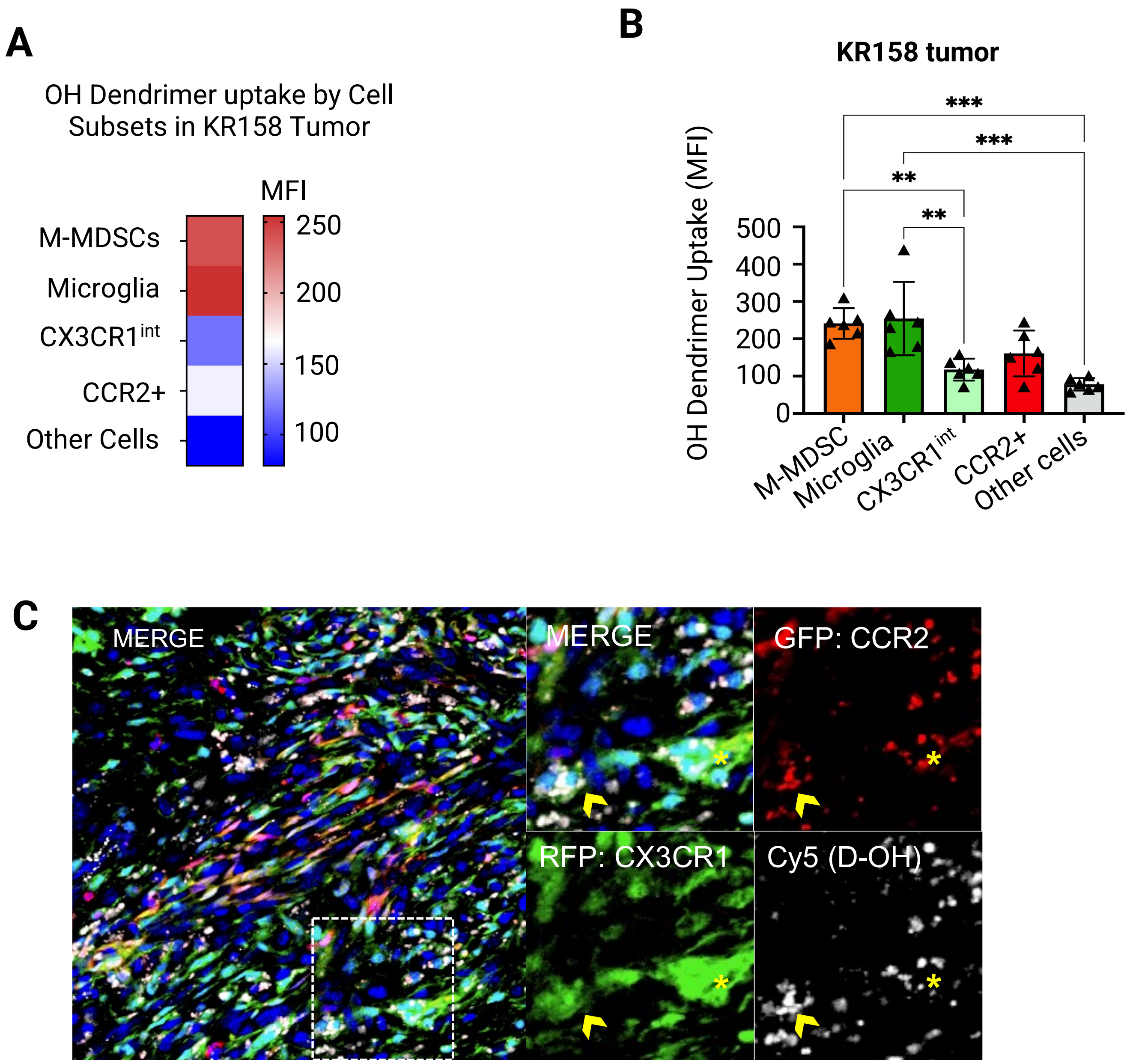


**Figure S1 A)**. The capacity of each cell subsets to uptake OH dendrimers within the KR158 gliomas at 24 hours post-injection. This is indicated by the MFI, which representatively measures the median number of dendrimers deposited per single cell. **B)** The statistical analysis of A). **C)** confocal microscopy image of dendrimer (white) distribution within the tumor (KR158) at 24 hours post-injection. Arrow and star indicate the intracellular localization of OH dendrimer with CCR2+/CX3CR1+ M-MDSCs. Red: RFP/CCR2; Green: GFP/CX3CR1; Blue: DAPI. White: dendrimer.

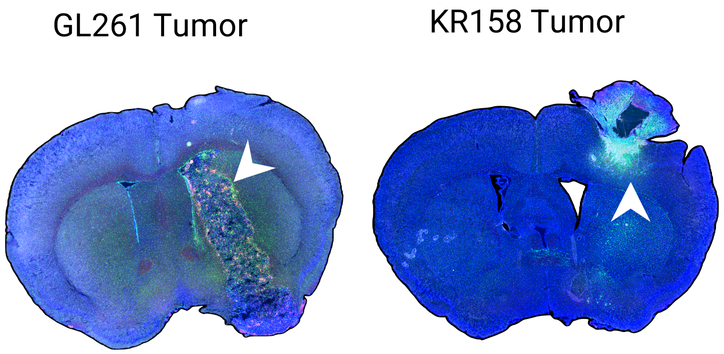


**Figure S2.** Representative fluorescence image of GL261 and KR158 tumor. Red: RFP/CCR2; Green: GFP/CX3CR1; Blue: DAPI. Arrow indicates the tumor stroma.

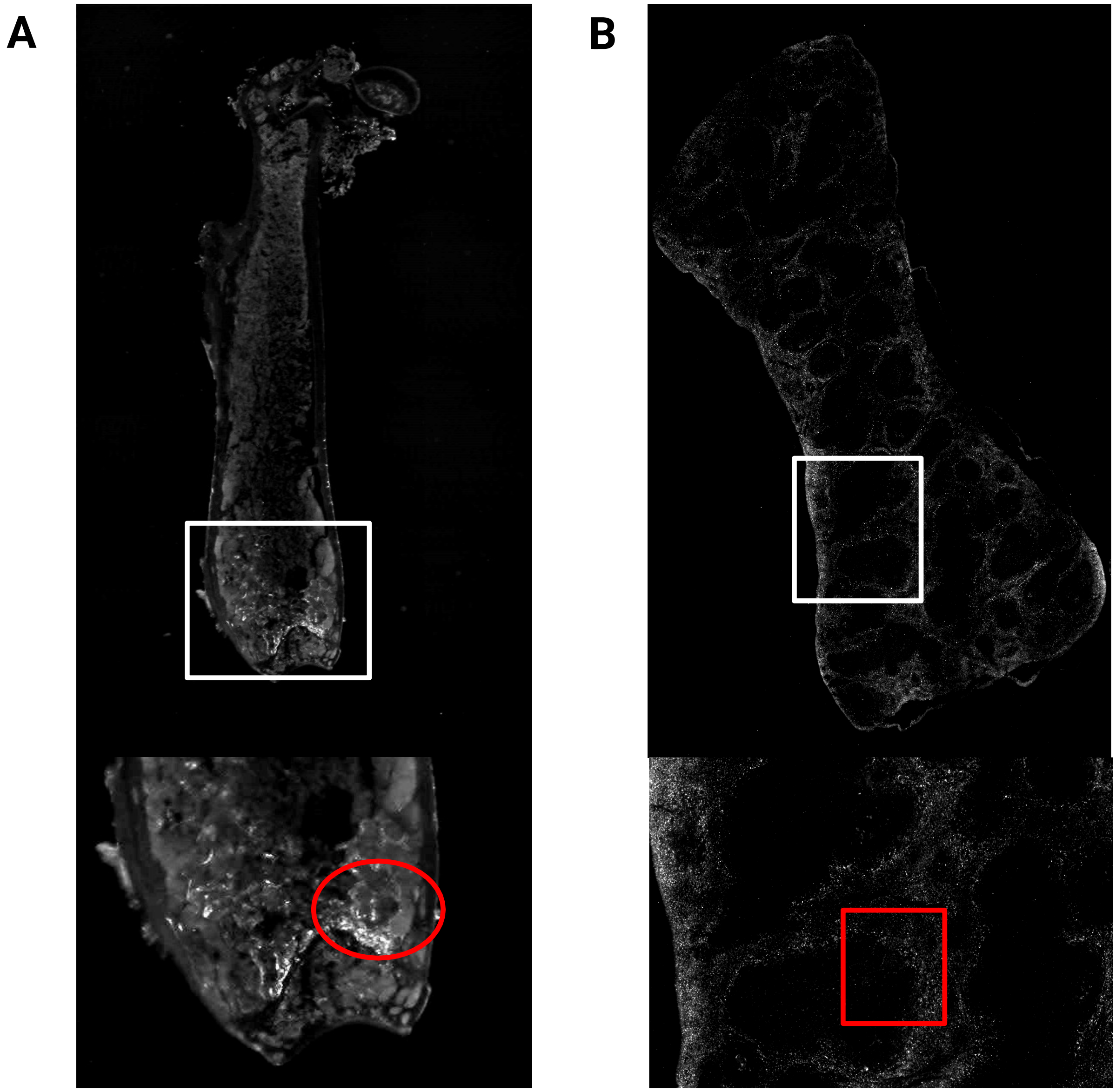


**Figure S3 A)** Murine bone marrow sagittal cross section showing OH dendrimer (Cy5) signal 24 hours post 50mg/kg injection of (OH) dendrimer. Image generated by fluorescence microscopy 4X magnification panel and optimized for brightness and contrast using Fiji. White box highlighting image zoom panel. Below showing OH dendrimer in the monocyte rich red marrow of spongy bone (red circle). **B)** Murine spleen sagittal cross section showing dendrimer (Cy5) signal 24 hours post 50mg/kg injection of OH dendrimer. Image generated by fluorescence microscopy 10X magnification panel. White box highlighting image zoom panel. Below showing dendrimer deposition in monocyte rich red pulp and exclusion from lymphocyte white pulp zones.

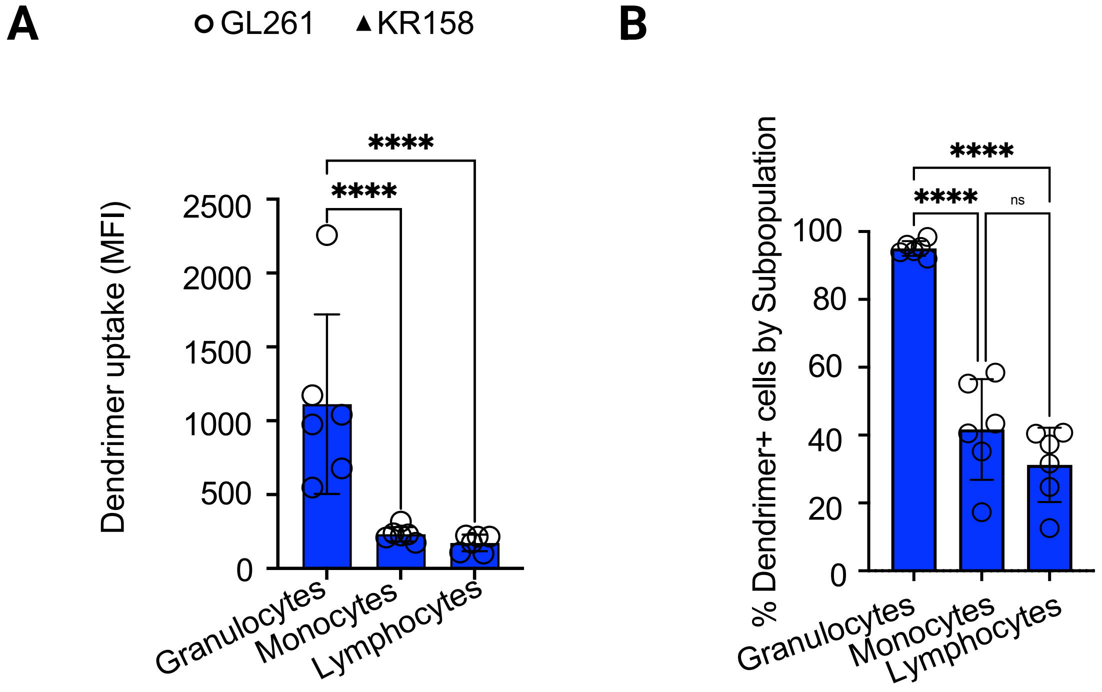


**Figure S4. Flow cytometry analyses of dendrimer uptake in blood leukocytes. A)** Comparison of OH dendrimer uptake (MFI) in leukocyte subpopulations of GL261 and KR158 tumor-bearing mice. n=6. **B)** Comparison of the percent of dendrimer positive leukocytes by subpopulations of GL261 and KR158 tumor-bearing mice. n=6. *p<0.05, **p<0.001, ***p=0.0001, ****p<0.0001

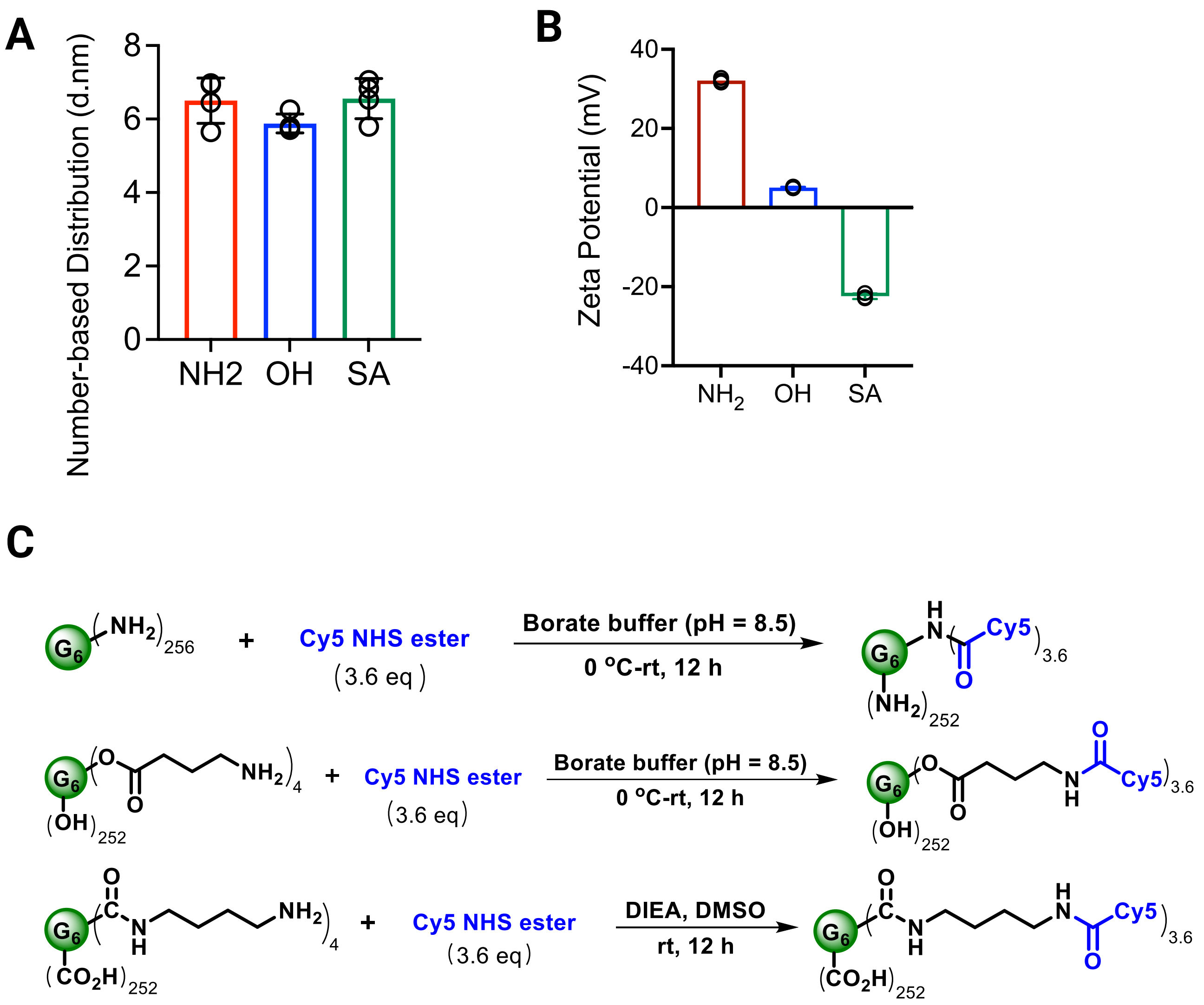


**Figure S5 A)** The number-average mean-based size and **B)** ζ-potential (mV) of hydroxyl (OH), succinamic acid (SA), and amine (NH2) dendrimers. The size and ζ-potential were measured by dynamic light scattering (DLS) and electrophoretic light scattering (ELS). **C)** Schematic illustration shows the labeling of Cy5 to NH2 (top), OH (middle), and SA (bottom) G6-PAMAM dendrimers. The Cy5 labelling of G6 PAMAM dendrimer (D-Cy5)1 and synthesis of hydroxyl surface bi-functional G6-PAMAM dendrimer and acid surface bi-functional G6-PAMAM dendrimer was carried out using a reported literature method2. The amine surface G6-PAMAM dendrimer (50 mg) was dissolved in borate buffer (2 mL, pH 8.5) at room temperature. The reaction mixture was cooled to 0 oC and Cy5 mono NHS ester (3.6 eq, 2.5 mg) in DMSO (1 mL) was added. The reaction mixture was allowed to stir overnight and lyophilized. The obtained crude product was dissolved in water and dialyzed (membrane cutoff = 12-14 kDa) against DI water for 24 h with successive change of water every 3 h. The collected dialysis bag water was lyophilized to get the amine surface D-Cy5 (48 mg). The Cy5 labelling of bi-functional hydroxyl surface PAMAM dendrimer was conducted by following the above procedure. The acid surface bi-functional PAMAM dendrimer (0.000590 mmol, 1.0 eq, 50 mg) was dissolved in DMSO (2 mL) at room temperature, to this solution was added Cy5 NHS ester (0.0022 mmol, 3.66 eq, 1.72 mg) and DIEA (0.0065 mmol, 11 eq, 1.5 uL). The resulting reaction mixture was allowed to stir overnight and dialyzed against DMSO for 8 h by changing the solvent at least two times, which was further dialyzed against DMF for 12 h by changing the solvent two times. The collected solvent was evaporated under high vacuum and the compound was dissolved in DI water, dialyzed against DI water for 6 h by changing the solvent every 2 hours. The collected water was lyophilized for 24 h to get pure acid surface D-Cy5 (45 mg).

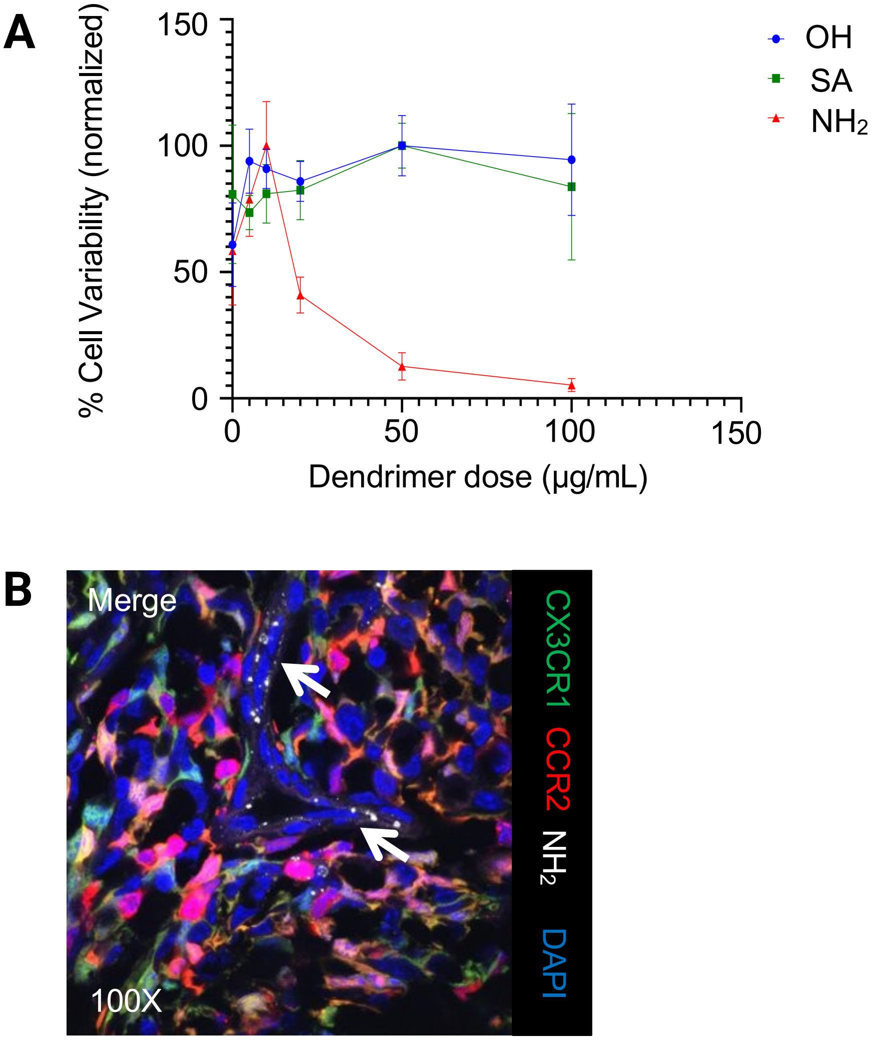


**Figure S6 A)** *In vitro* evaluation of the dose-dependent toxicity of NH2, OH, and SA dendrimers on primary M-MDSCs by cell-titer blue assay. Positive control: cells treated with 1% Triton-X, Negative control: cell without treatment. **B)** At 24 hours after systemic injection, NH2 dendrimers (10mg/kg) co-localized with the endothelial cell (indicated by arrow) in tumor stroma of a KR158 glioma established from $CCR2^{WT/RFP}CX3CR1^{WT/GFP}$ transgenic mouse.

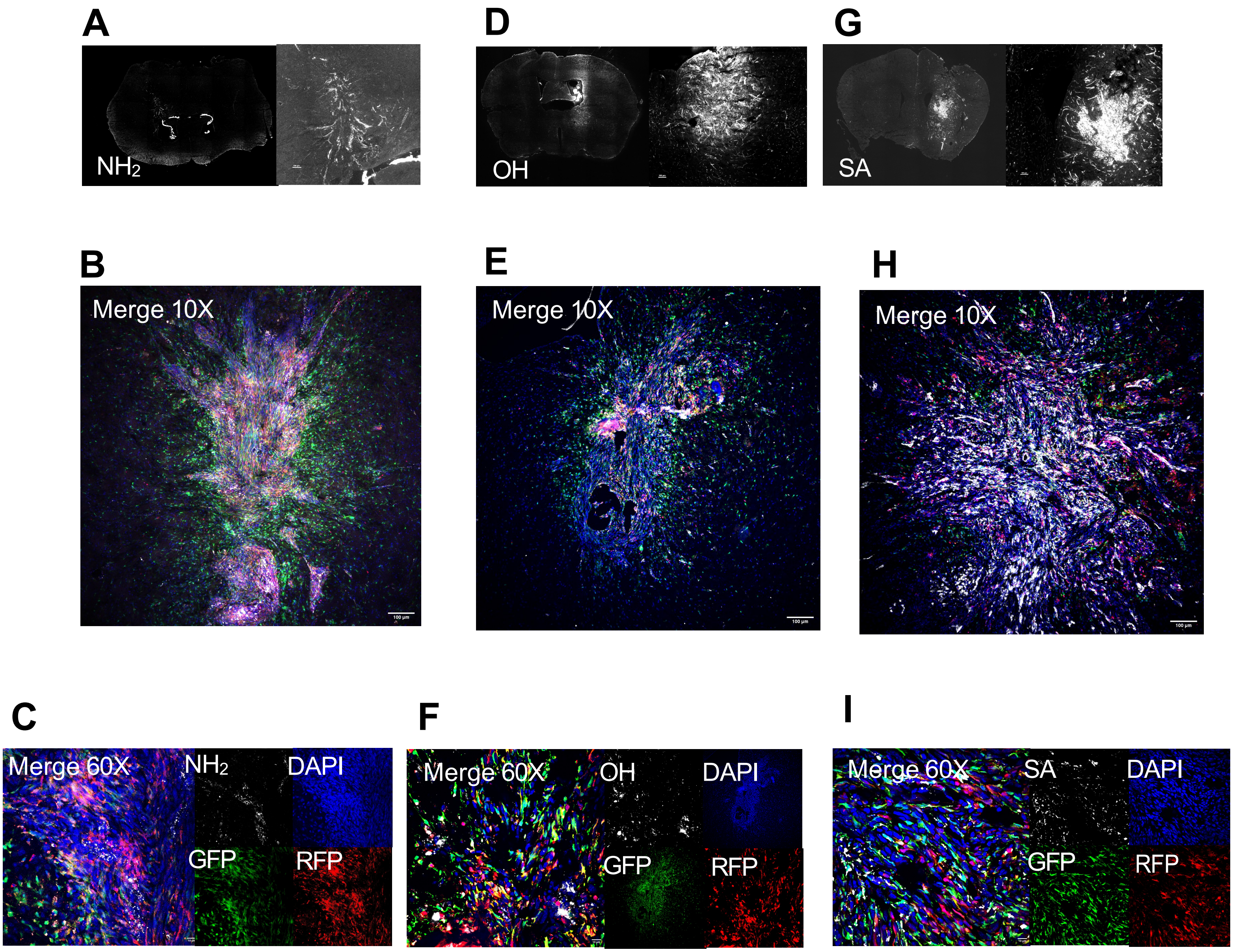


**Figure S7 A)** NH2 dendrimer distribution within the KR158 tumor at 24 hours post-injection (10mg/kg), Image based on the widefield fluorescence microscopy (left panel) with zoom (right panel). **B)** 10X merge confocal of tumor from A. **C)** 60X merge confocal for amine dendrimer tumor shown in panel B. **D)** OH dendrimer distribution within the KR158 tumor at 24 hours post-injection (50mg/kg), Image based on widefield fluorescence microscopy (left panel) with zoom (right panel). **E)** 10X merge confocal of tumor from tumor shown in panel D. **F)** 60X merge confocal for amine dendrimer tumor shown in panel E. **G)** SA dendrimer distribution within the KR158 tumor at 24 hours post-injection (50mg/kg), Image based on widefield fluorescence microscopy (left panel) with zoom (right panel). **H)** 10X merge confocal of tumor from tumor shown in panel G. **I)** 60X merge confocal for amine dendrimer tumor shown in panel H. Red = CCR2. Green = CX3CR1. White = Cy5-labeled dendrimer. Blue = DAPI (Nucleus).

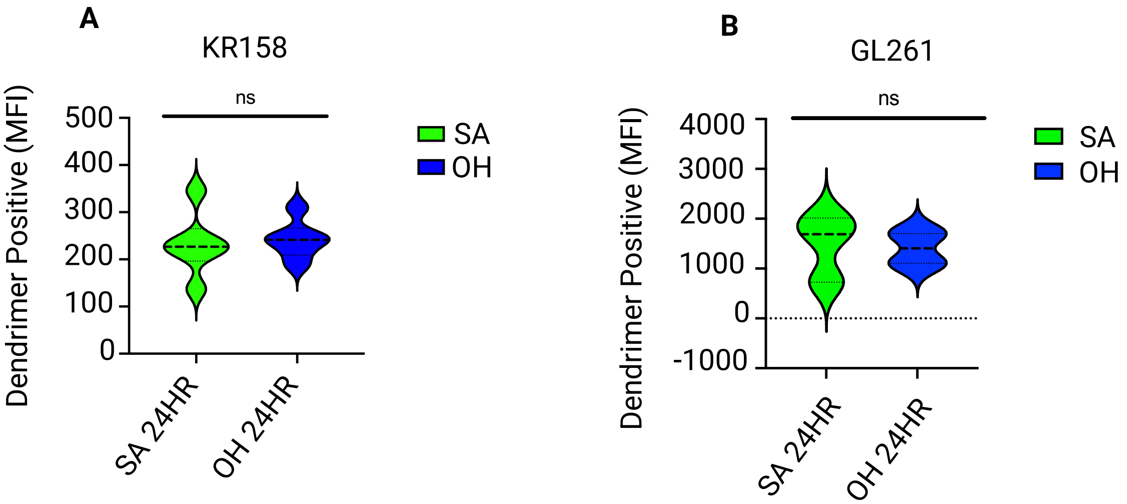


**Figure S8.** Comparison of overall tumor depositions of SA and OH dendrimers (indicated by Cy5 MFI) in KR158 tumor **A)** and GL261 tumor **B)**. NS: no statistical significance.

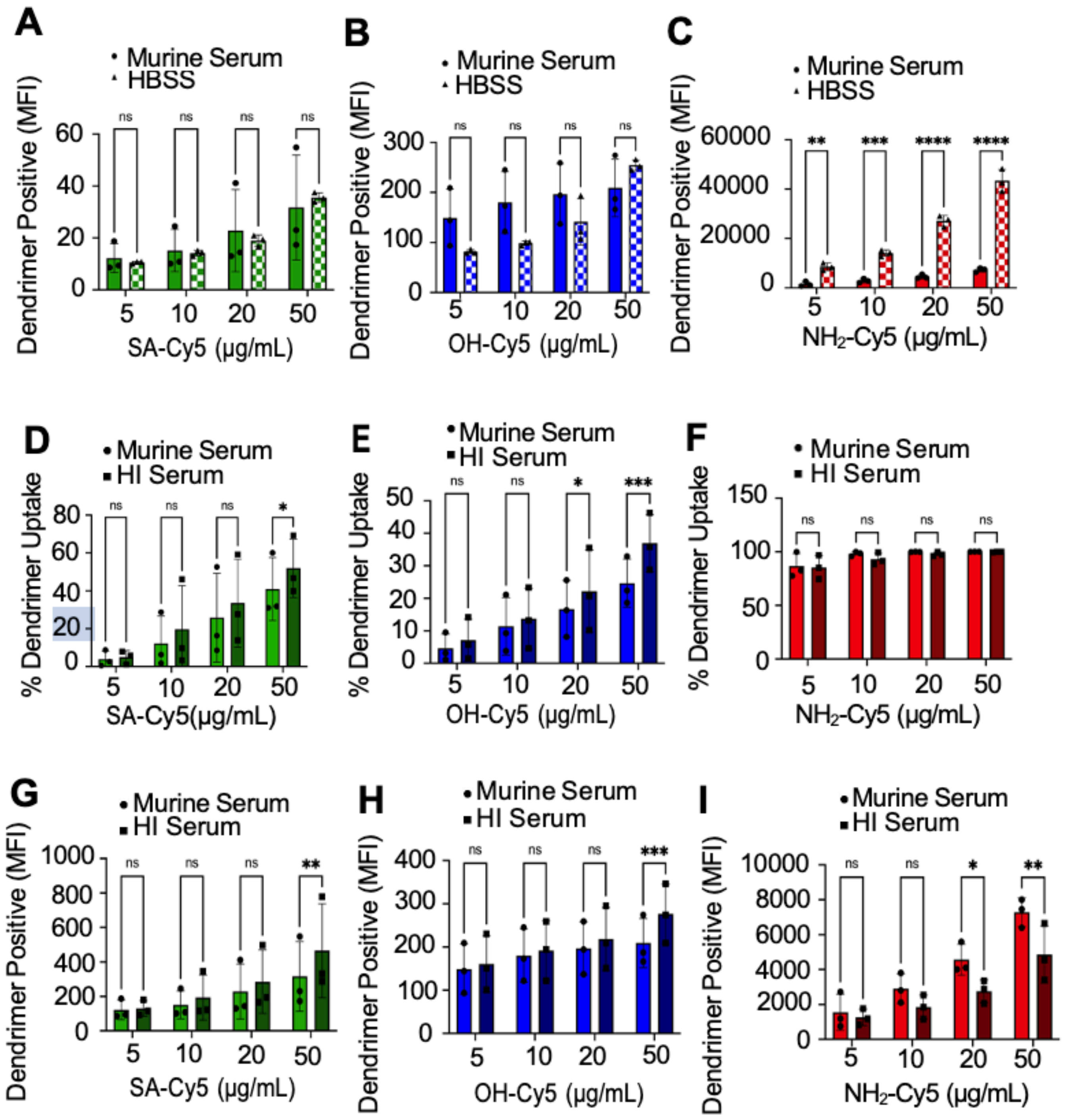


**Figure S9 A-C)** Dendrimer uptake by M-MDSCs comparing SA, OH, and $NH_2$ incubated in serum free buffer (1X HBSS) versus native murine serum in 1X HBSS by median fluorescence intensity (MFI) using flow cytometry. **D-F)** Percentage of dendrimer positive M-MDSCs comparing SA, OH, and $NH_2$ surface functionality with either native (murine serum) or denatured (HI serum) protein coronas with increasing concentration. n=3 biological repeats. **G-I)** Corresponding dendrimer uptake from samples D-F as determined by flow cytometry Cy5+ MFI. *p<0.0332, **p<0.0021,***p<0.0002, ****p<0.0001
